# Multiomics integrative analysis reveals antagonistic roles of CBX2 and CBX7 in metabolic reprogramming of breast cancer

**DOI:** 10.1101/2020.08.06.239129

**Authors:** Mohammad Askandar Iqbal, Shumaila Siddiqui, Asad Ur Rehman, Farid Ahmad Siddiqui, Prithvi Singh, Bhupender Kumar, Daman Saluja

## Abstract

Striking similarity exists between metabolic changes associated with embryogenesis and tumorigenesis. Chromobox proteins-CBX2/4/6/7/8, core components of canonical polycomb repressor complex 1 (cPRC1), play essential roles in embryonic development and aberrantly expressed in breast cancer. Understanding how altered CBX expression relates to metabolic reprogramming in breast cancer may reveal vulnerabilities of therapeutic pertinence. Using transcriptomic and metabolomic data from breast cancer patients (N>3000 combined), we performed pathway-based analysis and identified outstanding roles of CBX2 and CBX7 in positive and negative regulation of glucose metabolism, respectively. Genetic ablation experiments validated the contrasting roles of two isoforms in cancer metabolism and cell growth. Furthermore, we provide evidence for the role of mTORC1 signaling in mediating contrary effects of CBX2 and CBX7 on breast cancer metabolism. Underpinning the biological significance of metabolic roles, CBX2 and CBX7 were found to be the most up- and down-regulated isoforms, respectively, in breast tumors compared to normal tissues. Moreover, CBX2 and CBX7 expression (not other isoforms) correlated strongly, but oppositely, with breast tumor subtype aggressiveness and the proliferation markers. Consistently, genomic data also showed higher amplification frequency of CBX2, not CBX7, in breast tumors. Highlighting the clinical significance of findings, disease-specific survival and drug sensitivity analysis revealed that CBX2 and CBX7 predicted patient outcome and sensitivity to FDA-approved clinical drugs. In summary, this work identifies novel cross-talk between CBX2/7 and breast tumor metabolism, and the results presented may have implications in strategies targeting breast cancer.

## 1.1 Introduction

Breast cancer is a major health challenge with over 2 million cases diagnosed worldwide in 2018, second highest after lung cancer (1). In India, breast cancer has ranked number one in cancer-related deaths among women, surpassing cervical cancer (2). Although, substantial progress has been made in breast cancer treatment strategies to bring down morbidity and mortality; the patient outcome, particularly for aggressive breast cancers, remains poor. This necessitates the identification of the oncogenic mechanisms responsible for breast carcinogenesis and their evaluation for clinical relevance.

Nearly a century ago, Otto Warburg observed unusual conversion of glucose into lactate by cultured cancer cells even in the presence of ample oxygen, a phenomenon known as Warburg effect or aerobic glycolysis (3). Warburg effect along with other metabolic alterations essentially constitutes metabolic reprogramming, a key adaptation by cancer cells to support their rapid proliferation (4). Metabolic reprogramming discriminates tumor cells from their normal counterparts, thus, holding immense therapeutic significance (5). Aerobic glycolysis plays a central role in channeling glucose carbons for biomass production by branching-off pathways which rely on glycolytic intermediates as substrates, thus, prioritizing anabolism over catabolism (6). Elevated lactate production by cancer cells helps conserve glucose carbons for anabolic processes rather than ATP production via oxidative phosphorylation (7). Besides, glycolysis also serves as a source of rapid ATP production in cancer cells as it does in skeletal muscle during strenuous exercise (8). Notably, FDG-PET (^18^Fluorodeoxyglucose-positron emission tomography) exploits addiction of cancer cells to glucose for clinical imaging of primary and secondary tumors (9). It has now become increasingly evident that benefits of glycolysis extend beyond metabolism and role of altered glycolysis has been implicated in transcriptional regulation (10), epigenetic regulation (11, 12), immune-escape (13), cell cycle (14), mitotic spindle (15), metastasis (16), and inflammation (17). Moreover, aerobic glycolysis and associated pathways contribute to chemo- and radio-resistance in cancer (18–20). Combinatorial treatments with compounds inhibiting glycolysis have shown synergy in decreasing triple-negative breast cancer cell viability (21). Taken together, aerobic glycolysis is critical for tumor growth and survival, thus, it is important to elucidate the mechanisms that contribute to its regulation in breast cancer.

Metabolic changes play essential roles during embryogenesis and tumorigenesis (22, 23). For example, aerobic glycolysis facilitates biomass production during embryonic development (24). Moreover, metabolic changes are crucial in determining cellular fate and differentiation (25, 26). However, contrary to embryonic development where metabolic pathways are tightly regulated, cancer cells frequently acquire deregulation in metabolic pathways through mutations and epigenetic remodeling (27). Chromobox family members-CBX2, 4, 6, 7, and 8 (collectively referred as CBX, hereafter) are conserved components crucial for the activity of canonical polycomb repressor complex (cPRC1); and play a key role in embryonic development via transcriptional repression, necessary for maintaining cellular fate decisions (28, 29). CBXs are epigenetic readers which recruit PRC1 at specific methylated histones for transcriptional repression through chromatin compaction (30). Within PRC1 complex, CBX is reported to be mutually exclusive (31). CBX2 can also function independently of PRC1 complex (32). Deregulated CBX expression has been implicated in breast cancer (33–37). Recent evidence connects PRC1 with triggering oncogenic transcriptional programs in breast cancer (38). However, the relation between CBX with metabolic reprogramming remains unclear. With this background, we conjecture that altered CBX expression may play a role in breast cancer metabolic reprogramming.

Using the integrative approach, we attempt here to delineate the role of aberrant CBX expression in metabolic reprogramming and identify CBX2 and CBX7 (referred as CBX2/7, hereafter) as antagonistic regulators of aerobic glycolysis in breast cancer. Further, we evaluate the biological and clinical relevance of identified metabolic roles of CBX2 and CBX7 to show that these two isoforms are most differentially expressed in breast tumors and are informative about prognosis and drug sensitivity.

## 1.2 Results

### 1.2.1 Transcriptomic and metabolomic data are mutually corroborative to suggest opposing roles of CBX2 and CBX7 in breast cancer metabolism

To understand the role of CBX members in breast cancer metabolism, we queried transcriptomic data of tumor and normal tissue samples from clinically annotated METABRIC and TCGA datasets. As aerobic glycolysis is central to metabolic reprogramming, expression of glycolysis geneset (from Recon 1)(39) was analyzed. For a meaningful interpretation of gene expression information, we employed Pathifier (40) to assign a pathway deregulation score (PDS) to each tumor sample based on the deviation (in geneset expression) from normal sample. As shown in Fig. 1A, breast tumors exhibit highly deregulated glycolysis compared to normal tissue. To evaluate the association between CBX isoforms and glycolysis, a correlation was calculated between each CBX and glycolysis PDS across all samples. Strikingly, CBX2 and CBX7 stood-out in their positive and negative correlation, respectively, with glycolysis PDS (Fig. 1B). CBX4 and CBX8 correlation with glycolysis didn’t reproduce in two datasets. Although CBX6 correlation with glycolysis was consistent in datasets, CBX7 outperformed CBX6 in the strength of correlation with glycolysis (Fig. 1B, see related Fig. S1A). Further, samples were designated as CBX2High/Low or CBX7High/Low based on above (High) or below (Low) mean expression. CBX2High and CBX7Low samples showed higher glycolysis PDS compared to CBX2Low and CBX7High samples, respectively (Fig. 1C). Interestingly, glycolysis deregulation correlated with subtype aggressiveness with lowest and highest PDS in lumA and basal samples, respectively (Fig. 1D). To further substantiate these observations, we accessed breast tumor metabolomics data published by Terunuma et al. (41). Expectedly, metabolomics data showed up-regulation of glycolysis metabolites in tumor samples compared to normal tissues (Fig. 1E). Remarkably, the correlation of CBX2, 6 and 7 with glycolytic metabolites showed similarity to their correlation with glycolysis PDS (compare Fig. 1B and Fig. 1F). Likewise, glycolytic metabolites were found to be up-regulated in CBX2High and CBX7Low samples (Fig. 1G). Moreover, key metabolites of biosynthetic pathways that branch-off from glycolysis were also found to be up-regulated in CBX2High and CBX7Low tumor samples (Fig. S1B). Overall, these data demonstrate agreement between gene expression and metabolite data to indicate conflicting roles of CBX2/7 in breast cancer metabolism.

**Figure 1:**
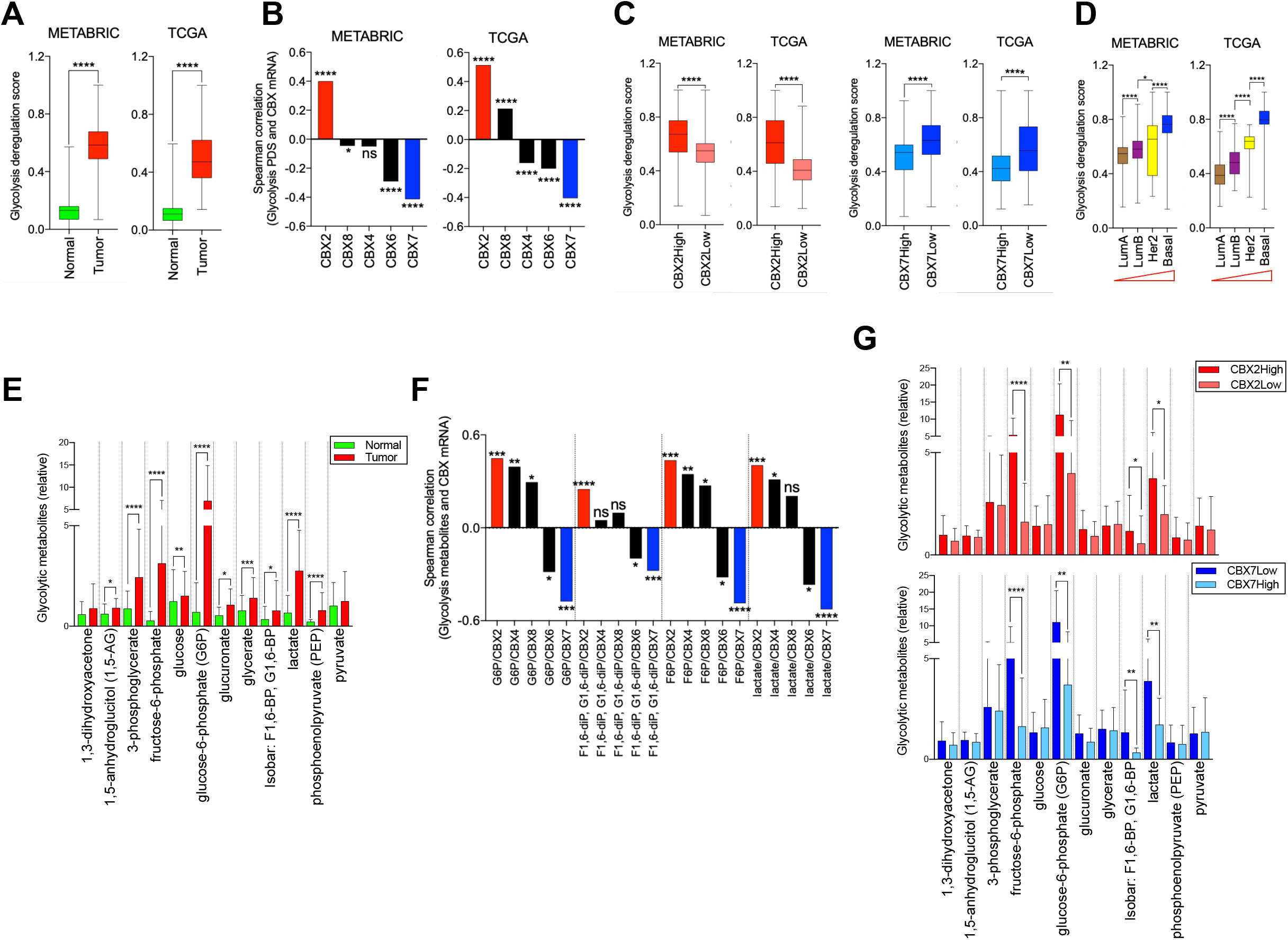
Analysis of transcriptomic and metabolomic data from breast tumors. A) Glycolysis is highly deregulated in breast tumors of METABRIC (N=1992) and TCGA (N=1104) compared to normal tissues (N=144 and N=102 in METABRIC and TCGA, respectively). Gene-expression based deregulation score for normal and tumors was calculated using Pathifier tool (see text). B) Spearman correlation between glycolysis deregulation score (PDS) and mRNA expression of CBX members in breast tumors of METABRIC and TCGA. C) CBX2High (N=815) and CBX7Low (1060) tumor samples showed higher glycolysis PDS compared to their counterparts, consistent with their oncogenic and tumor-suppressive roles, respectively. D) Glycolysis deregulation score increased from lumA to basal samples, indicating an association of deregulated glycolysis with subtype aggressiveness. The red triangles indicate an increase in aggressiveness from left to right. E) Metabolomics data from Terunuma et al. showing significantly higher intracellular levels of glycolysis intermediates in breast tumors compared to normal tissues (Normals=65, Tumors=67). F) Spearman correlation analysis between key glycolytic metabolites and CBX mRNA in breast tumors from Terunuma et al. G) Breast tumors (Terunuma et al.) were divided as CBX2High/Low and CBX7High/Low and glycolytic metabolite abundance was compared to show significant differences. Box and whiskers plot represent minimum, maximum and median. *P* values were calculated using Kruskal-Wallis test or *t*-test and represented as \**P*<0.03, \*\**P*<0.0021, \*\*\**P* <0.0002, \*\*\*\**P*<0.0001.

### 1.2.2 Silencing of CBX2 and CBX7 reveal inverse effects on glycolysis, ATP production, viability, proliferation and biomass production

Transcriptomic and metabolomic results were concordant to implicate CBX2/7 in aerobic glycolysis. To experimentally corroborate, we silenced CBX2 or CBX7 in MDA-MB-231 and MCF7 cells using siRNA approach; followed by measurements of glucose uptake, lactate release, glycolytic ATP production, viability, proliferation and biomass production. To ensure measurements of only glycolytic ATP, all experimental cells were given background treatment with 1 μM oligomycin (mitochondrial ATP synthase inhibitor). In agreement with patient tumor data, silencing of CBX2 and CBX7 showed opposite effects on all six measured end-points (Fig. 2A-F).

**Figure 2:**
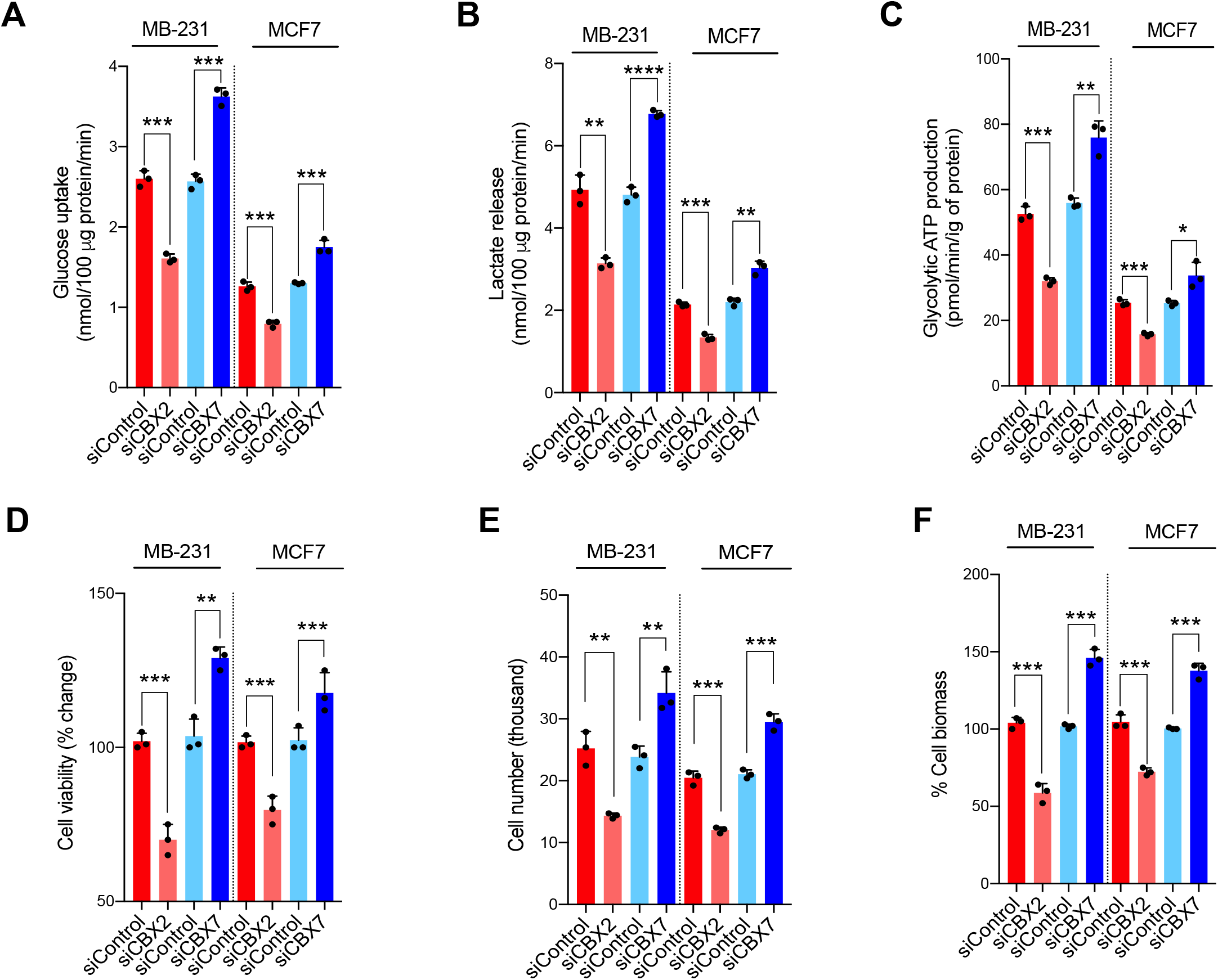
Effect of CBX2 and CBX7 silencing on glucose uptake, lactate release, glycolytic ATP production, cell growth and biomass production. A-C) MDA-MB-231 cells were either transfected with siControl (non-targeting pool siRNA) or siCBX2 or siCBX7 (SMARTpool siRNA) and after 48 hours media was collected to assess glucose uptake and lactate release using commercial kits as described in materials and methods section. Intracellular ATP was extracted from siControl, siCBX2 and siCBX7 transfected cells after 48 hours, as described in materials and methods. D-F) siControl, siCBX2 and siCBX7 transfected cells were assessed for cell viability, proliferation and biomass production. Bars represent mean ± SD from 3 independent biological replicates with readings taken in triplicate. *t*-test was used to calculate *P* values represented as \*\**P*<0.0021, \*\*\**P* <0.0002, \*\*\*\**P*<0.0001.

CBX7 over-expression mirrored the effects of CBX2 silencing (Fig. S2A). Consistent with effect of CBX2/7 silencing on proliferation (Fig. 2E), CBX2/7 protein correlated significantly with proliferation markers CCNB1 and Ki67 in TCGA breast tumor tissues (Fig. S2B-C). To further establish that changes in viability, proliferation and biomass are indeed because of the decreased glycolysis induced upon CBX2 and CBX7 silencing, MDA-MB-231 cells were treated with 2-deoxyglucose (2DG), a glucose analog that competes with glucose and blocks the conversion of glucose to glucose-6-phosphate, thus inhibiting glycolysis (42). Reduction in viability, proliferation and biomass production upon 2DG treatment demonstrated the role of glycolysis in controlling cell growth (Fig. S3). Moreover, CBX2/7 silencing resulted in changes in glycolysis gene signature (Fig. S3D). Overall, these *in vitro* results were found to be consistent with the findings of the transcripto-metabolomic analysis and validated the roles of CBX2 and CBX7 in metabolic reprogramming of breast cancer.

### 1.2.3 Evidence for the role of mTORC1 signaling

Dysregulated signaling is a common alteration in cancers and has been linked to abnormal nutrient uptake and metabolism in cancer (43). Oncogenic mutations are a frequent occurrence in receptors and/or regulators of growth signaling, an important requirement by cancer cells for unchecked nutrient-uptake (e.g. glucose) and altered metabolism (44). We conjectured that CBX2 and CBX7 may modulate growth signaling to induce metabolic reprogramming in breast tumors. To test the hypothesis, genesets of hallmark cancer signaling pathways were retrieved from MSigDB by Broad Institute (45) and subjected to Pathifier for calculation of deregulation scores of all hallmark signaling pathways in all samples of METABRIC and TCGA datasets. Correlation analysis of PDS of signaling hallmarks with CBX2/7 revealed the strongest correlation of both isoforms with mTORC1 signaling in either direction, reproduced across both datasets (Fig. 3A; also see Fig. S4 for correlation plots). Moreover, CBX2/7 protein correlated in opposite directions with phosphorylation of ribosomal S6 protein at serine 235/236 and serine 240/244 in TCGA tumor samples, further implying the role of mTORC1 signaling (Fig. 3B). Importantly, phosphorylation of ribosomal S6 protein is considered as a conserved marker of mTOR signaling state (46). Next, for more direct evidence, CBX2 and CBX7 were silenced in MDA-MB-231 and MCF7 cells using SMARTpool siRNA (combines four gene-specific siRNA for effective gene silencing) followed by immunoblotting to detect phosphorylation of ribosomal S6 protein. As shown in Fig 3C; CBX2 or CBX7 silencing decreased or increased phosphorylation of ribosomal S6 protein, respectively. Quantification of blots is provided in Fig. S5A. Further, reduction of glycolysis in MDA-MB-231 and MCF7 upon treatment with standard mTORC1 signaling inhibitor rapamycin validated the role of mTORC1 signaling in regulation of glycolysis (Fig. S5). Together, these results hint at the role of CBX2 and CBX7 in modulating mTORC1 pathway to promote aerobic glycolysis in breast cancer.

**Figure 3:**
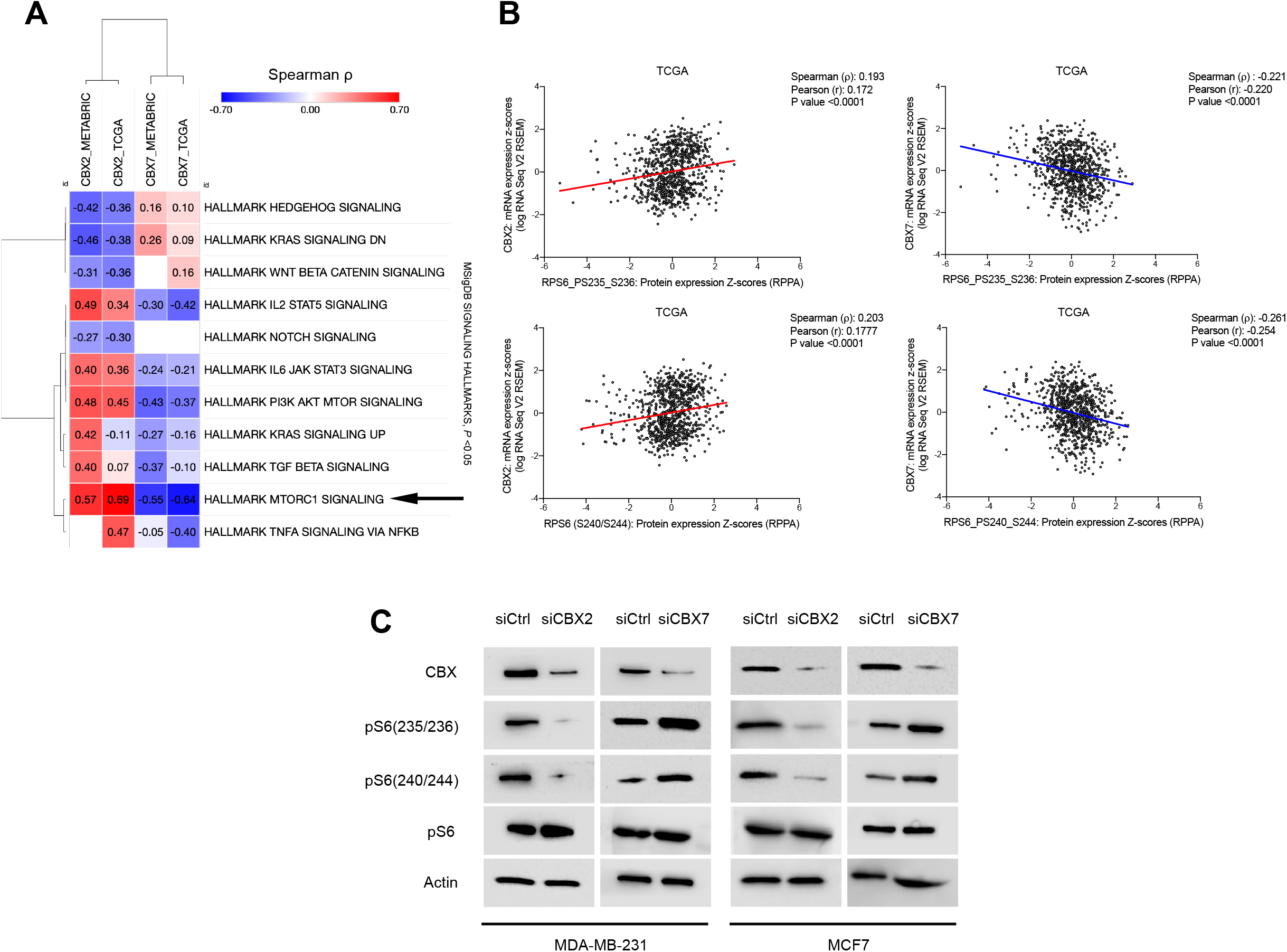
Effect of CBX2/7 silencing on mTORC1 signaling. A) Heatmap showing correlations of CBX2 and CBX7 mRNA with deregulation scores (calculated using Pathifier as described in methods section) of MSigDB signaling hallmarks of cancer. White squares in heatmap represents non-significant correlation. B) Correlations of CBX2 and CBX7 mRNA with ribosomal phosphoS6 (S235/236 and S240/244) protein expression in TCGA breast tumors (see text). C) Immunoblots showing inhibition of mTORC1 signaling upon CBX2 silencing while activation in CBX7 silenced MDA-MB-231 and MCF7 cells.

### 1.2.4 CBX2 and CBX7 are the most differentially expressed isoforms compared to normal tissue and exhibit opposing correlation with breast cancer aggressiveness

Discriminatory expression in tumors compared to normal and over-expression in aggressive tumors are important indices that define the significance of a gene in cancer. To find out if CBX2 and CBX7 meet these criteria; and to further validate the biological relevance of metabolic roles of CBX2/7, we performed the geno-transcriptomic analysis in METABRIC and TCGA datasets. Reproducibly in both datasets, CBX2 was found to be the most up-regulated and CBX7 as the most down-regulated isoforms in breast tumors compared to normal tissues (Fig. 4A-B). Notably, only CBX2/7 correlated with subtype aggressiveness, and not others, a striking similarity to the correlation of glycolysis deregulation with breast cancer aggressiveness (Fig. 4C and Fig. 1D). As glycolysis and tumor aggressiveness relates to high rates of proliferation (13, 47), CBX2/7 protein (not CBX4/6/8) correlated significantly with protein levels of tumor proliferation marker Ki67 in TCGA breast tumors (Fig. S2). Copy number analysis showed higher amplification frequency of CBX2 gene and almost negligible amplification of CBX7, suggesting oncogenic and tumor suppressive roles, respectively (Fig. 4D). Amplification of CBX2 resulted in increased mRNA but no change in mRNA of CBX7, in agreement with copy number data (Fig. 4D). With regard to regulation of gene expression, DNA methylation of levels CBX2/7 gene inversely and significantly correlated with CBX2 and CBX7 mRNA in tumor samples, suggesting the role of CpG methylation in regulation of their gene expression (Fig. S6A). Moreover, CBX2/7 mRNA correlated significantly with protein levels in TCGA breast tumor samples (Fig. S6B). CBX2 and CBX7 are over- and under-expressed in a subset of tumor samples; and percentage of CBX2 over- and CBX7 underexpressing samples increased with breast tumor aggressiveness in both datasets (Fig. 4F). In conclusion, these data highlighted CBX2/7 as the important players, particularly in aggressive breast cancer, an observation concurring with their metabolic functions.

**Figure 4:**
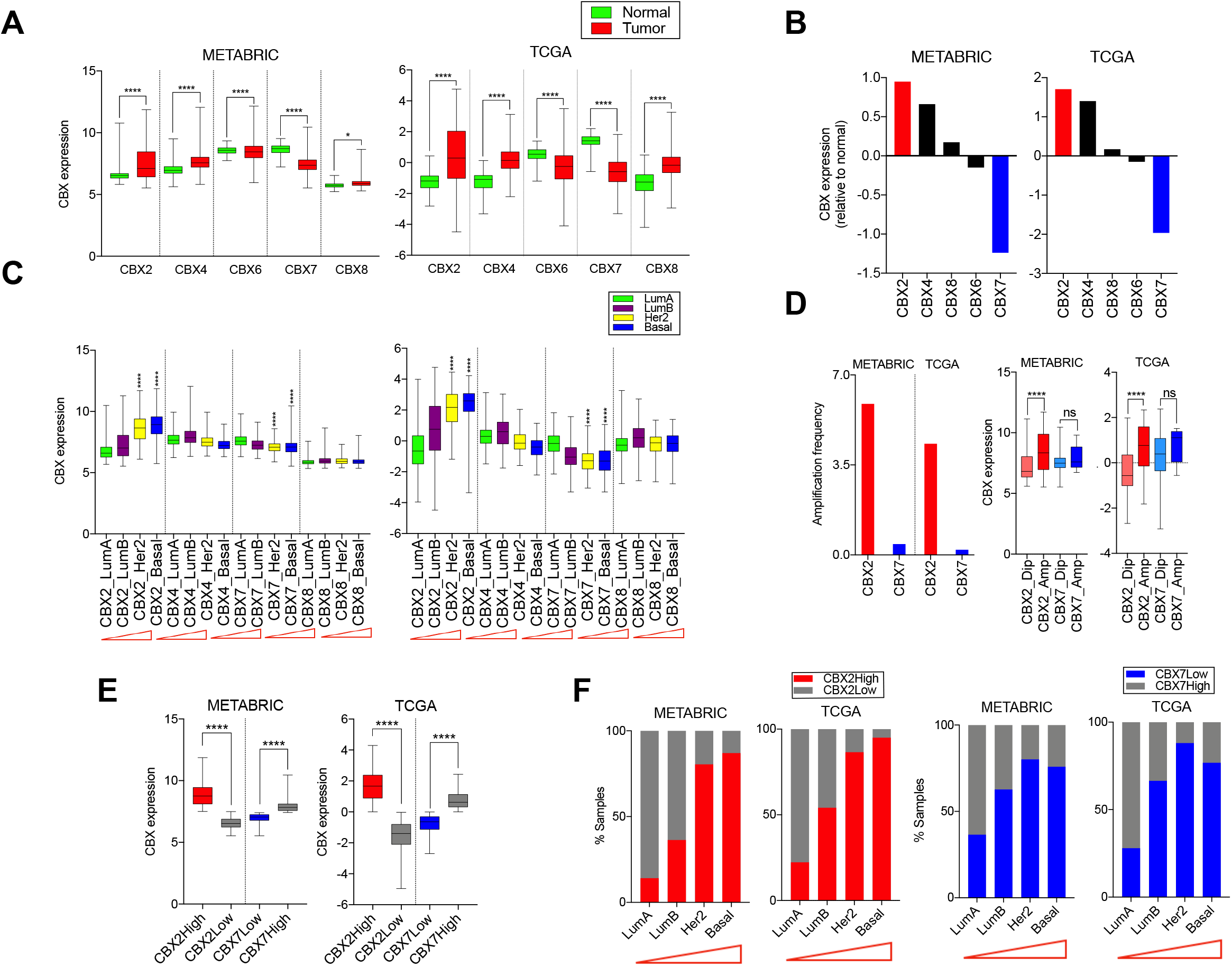
Genotranscriptomic analysis of CBX members in breast tumors and normal tissues. A) Expression of different CBX members in normal and tumor samples of METABRIC and TCGA cohorts. B) The highest and lowest expression of CBX2 and CBX7 respectively in breast tumors compared to normal, across both datasets. C) CBX2 and CBX7 were the only isoforms associated with subtype aggressiveness, as the CBX2 expression increases and CBX7 expression decreases from least aggressive lumA to most aggressive basal type tumors. The red triangle indicates an increase in aggressiveness from left to right. D) Copy number analysis shows higher amplification of CBX2 and almost no amplification of CBX7 in breast tumors of METABRIC and TCGA. Amplification resulted in increased mRNA of CBX2 but no change in CBX7 mRNA compared to diploid breast tumors of METABRIC and TCGA. E) CBX2 is over-expressed in a subset of breast tumors in both datasets (N=815/517 in METABRIC/TCGA). Likewise, CBX7 is under-expressed in a subset of tumors in both datasets (N=1060/545 in METABRIC/TCGA). F) Percentage of CBX2High and CBX7Low tumors increase with subtype aggressiveness. *P* values were calculated using the Kruskal-Wallis test or *t*-test and represented as \**P*<0.03 and \*\*\*\**P*<0.0001.

### 1.2.5 Expression of CBX2 and CBX7 is predictive of prognosis and sensitivity to anti-cancer drugs

To assess the clinical relevance of the antagonistic roles of CBX2 and CBX7 in modulating aerobic glycolysis, Kaplan-Meier survival plots were generated. Patient prognosis is a key clinical indicator of cancer aggressiveness. CBX2 over-expressing tumors exhibited poor disease-specific survival (DSS) with log-rank HR: 1.816 and *P* value <0.0001 (METABRIC) and log-rank HR: 1.902 and *P* value: 0.0039 (TCGA), Fig. 5A. On the contrary, lower CBX7 expression associated with worse prognosis with log-rank *P* value <0.0001 and HR: 1.866 (METABRIC) and log-rank *P* value: 0.0031 and HR: 2.098 (TCGA), Fig. 5A. Since CBX2High and CBX7Low tumors accumulated in basal/her2 subtype, which are typically more aggressive than other subtypes, the difference in prognosis may be due to basal/Her2 and not CBX2High and CBX7Low status of tumors. To rule out, we removed basal/her2 samples and examined if CBX2 and CBX7 expression still predict patient survival. CBX2High and CBX7Low predicted poor prognosis even after removal of basal and her2 subtype samples, demonstrating the prognostic relevance of two isoforms (Fig. 5B). However, CBX2/7 couldn’t predict survival within basal and her2 subtypes, reproducibly in both datasets (Fig. S7A-B). Also, other CBX members could not significantly and reproducibly predict patient survival (Fig. S7C). Further, tumor samples with higher glycolysis deregulation score exhibited poor survival compared to samples with lower deregulation score (Fig. S7D).

**Figure 5:**
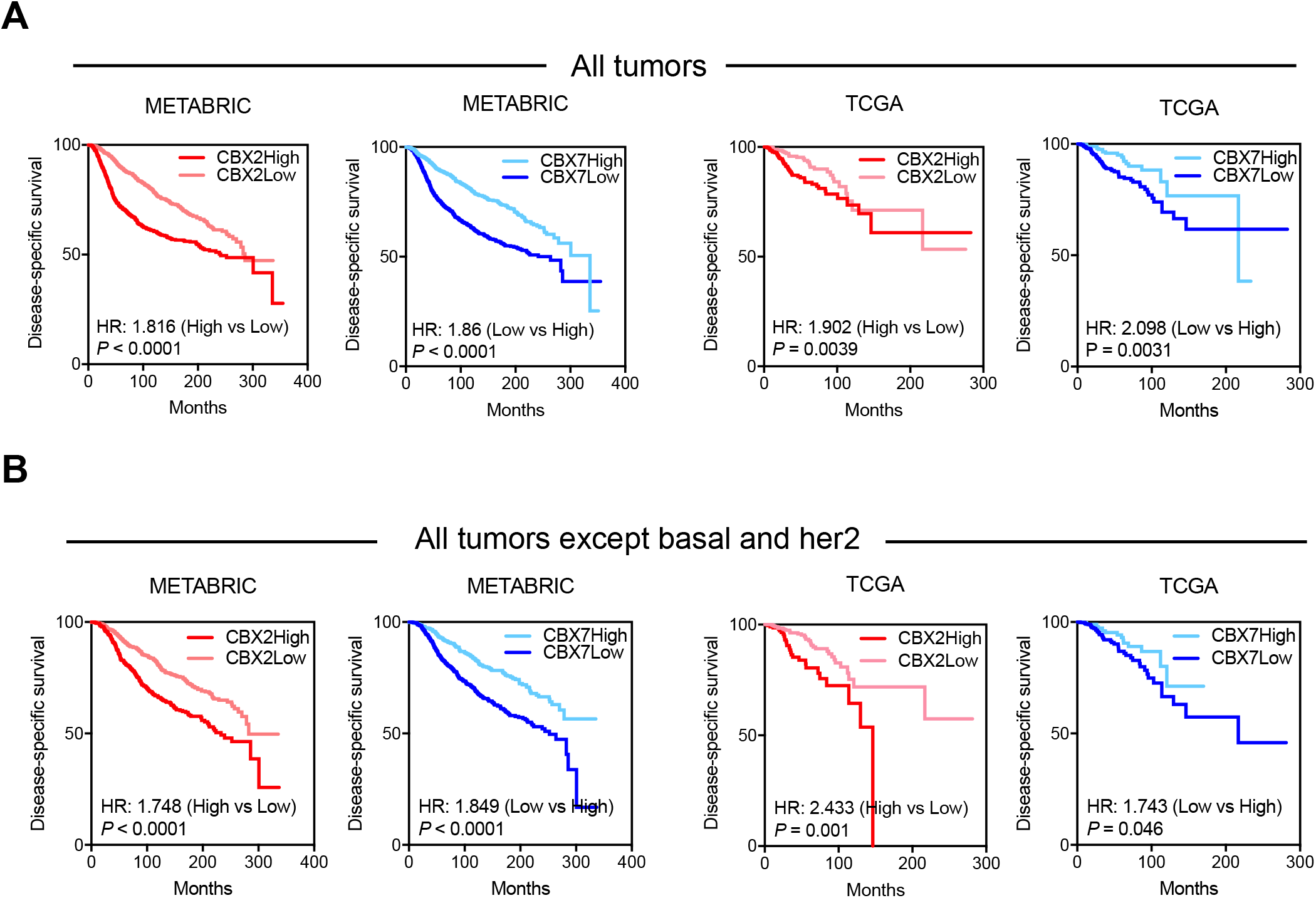
CBX2 and CBX7 expression informs of patient outcome. Kaplan-Meier survival curves showing A) CBX2High and CBX7Low correlate with poor clinical outcome in METABRIC (N=1986) and TCGA (N=1082) breast tumors. B) CBX2High and CBX7Low predicted poor prognosis in METABRIC (N=1415) and TCGA (N=876) tumors excluding basal and her2 subtype.

Chemotherapeutics is the mainstay treatment option for the majority of cancers. However, not all patients respond uniformly to a particular drug, a major hurdle in the clinical management of cancer. Identification of genes that could predict tumor response is critical for successful chemotherapeutics. Therefore, to further evaluate the clinical relevance, we investigated the role of oncogenic CBX2 in determining sensitivity to anti-cancer drugs. For this purpose, we accessed drug sensitivity data of 77 FDA-approved and investigational therapeutic compounds in a panel of 49 breast cancer cell lines along with expression data from Heiser et al. (48). We found that 16 compounds showed significantly different sensitivities in CBX2High and CBX2Low cell lines, as depicted in heatmap based on GI50 values (Fig. 6A). GI50 is the concentration needed for each drug to inhibit proliferation by 50%, where GI indicates growth inhibition (48). Of note, CBX2High cell lines were relatively sensitive to 4- and resistant to 12 out of 16 drugs and vice-versa for CBX2Low cell lines, suggesting that CBX2 mRNA could inform about a cell line’s response as relatively resistant or sensitive to these drugs (Fig. 6A). Interestingly, CBX2High cell lines were found to be sensitive to rapamycin, consistent with our observation that CBX2 modulates mTORC1 signaling (Fig. 6B). Moreover, CBX2High cells were sensitive to methotrexate (Fig. 6B), a chemotherapeutic drug used in treatment of breast cancers (49). However, effect of CBX7 on mTORC1 signaling did not correlate in terms of sensitivity to rapamycin as reflected in a [Fig. S8]. This could be attributed to the known complexity of rapamycin sensitivity in breast cancer cell lines (50, 51). For instance, PTEN, a critical regulator of Akt/mTOR pathway couldn’t predict sensitivity to rapamycin (51). Moreover, we did not observe overall antagonism in sensitivities to drugs in CBX2/7High cell lines, indicating that glycolysis/mTORC1 may not be the only determinants of drug sensitivity. Rather, overall functions of CBX2/7 (not necessarily antagonistic) may be defining sensitivities of breast cancer cell lines. Regardless, these results provide evidence of the relevance of CBX2/7 in predicting sensitivity to anti-cancer drugs and suggest that CBX2High breast tumors may be more likely to benefit from methotrexate and rapamycin treatment.

**Figure 6:**
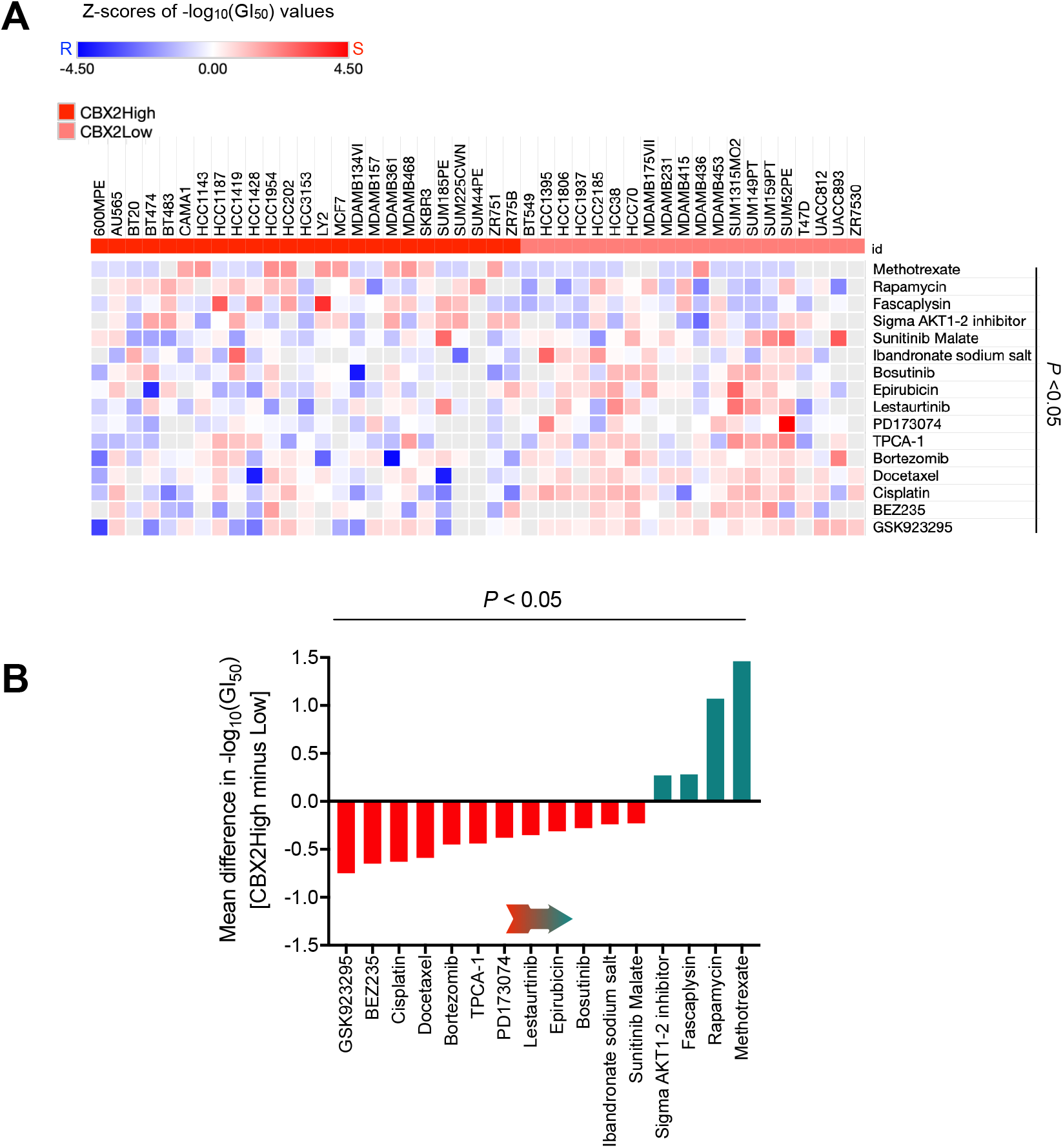
CBX2 expression and sensitivity to anti-cancer drugs. A) Heatmap showing differential sensitivity of CBX2High and CBX2Low cell lines to 12 anti-cancer drugs (FDA-approved or in preclinical stage) for treatment of breast cancer. CBX2High cell lines were found to be sensitive to 9 out of 12 drugs. Light grey squares represent value not available. B) Bar graph showing the mean difference in sensitivities of all 16 drugs. CBX2High cell lines were most sensitive to methotrexate and rapamycin. Red bars indicate drugs to which CBX2High cell lines are relatively resistant and green bars indicate drugs with higher sensitivity for CBX2High cell lines. Arrow shows the direction of increasing sensitivity.

## 1.3 Discussion

Metabolic transformation plays an essential role in supporting tumor progression. It is now well-appreciated that cancer glycolysis doesn’t just satisfy metabolic needs but also contributes to other hallmark properties of cancer (52). Accordingly, the Warburg effect plays a role in the promotion of cancer aggressiveness and drug resistance (53). Therefore, it is important to unravel the oncogenic mechanisms responsible for revving up glycolysis in cancer. Moreover, elucidating the clinical relevance of such mechanisms for their exploitation in anti-cancer strategies is required. In this work, we delineate the conflicting roles of CBX2/7 in the regulation of glycolysis in breast cancer and establish their clinical relevance.

Tumors addicted to glucose exhibit escalated rates of glycolysis, as indicated by increased intracellular concentrations of glycolytic metabolites in tumors compared to normal tissues (Fig.). A large majority of glucose that enters tumor cell gets quickly phosphorylated by hexokinase to prevent its escape (54), that’s why G6P essentially represents glucose taken-up by tumor cells. This explains why G6P is the most abundant glycolytic metabolite in tumor cells compared to normal (Fig. 1E). Lower abundance of F6P (compared to G6P) indicates that not all G6P enters glycolysis, and some G6P enters the pentose phosphate pathway (PPP) which branches-off from glycolysis (55). Lower levels of F1,6diP (compared to F6P) may be due to utilization of F6P into the synthesis of amino sugars via hexosamine biosynthesis pathway (HBP) (56). Use of glucose carbons into PPP and HBP, suggests that a large chunk of glucose taken-up by tumor cells is channelled for anabolic synthesis. Metabolite data confirmed the up-regulated levels of key metabolites of PPP and HBP pathway in CBX2High and CBX7Low tumors, thus, reflecting on the biology of glycolysis and shunting pathways (Fig. S1B). *In vitro* data showing the changes in glycolysis and cell growth upon CBX2/7 silencing complemented the metabolomics data to show differential glycolytic roles of CBX2 and CBX7. For instance, increased levels of PPP and HBP metabolites in CBX2High tumors were consistent with decreased biomass in siCBX2 cells. Moreover, a drop in viability and proliferation of siCBX2 cells reflects on the dependency of MDA-MB-231 cells on biomass production using glycolytic flux and vice-versa for CBX7. Overall, transcripto-metabolomics data of tumor samples along with experimental data conclusively demonstrate the role of CBX2/7 in glycolytic regulation. Strong concurrence between transcriptomic and metabolomic data from breast tumors not only highlight the robustness of our observations; but also signify that the gene expression pattern could predict metabolic behaviour in breast cancer patients (Fig. 1). CBX2 and CBX7 as the most differentially expressed isoforms in breast tumors compared to normal and the only two members whose expression is associated with breast cancer aggressiveness highlights the biological relevance of their determined metabolic roles (Fig. 4). Considering the complex and heterogeneous nature of glycolytic regulation concerning the players and mechanism involved (57–60), unraveling the mTORC1-mediated metabolic roles CBX2 and CBX7 may improve our understanding about how glucose metabolism is regulated in breast cancer. mTORC1 signaling is a known determinant of cancer metabolism and frequently deregulated in breast cancer (61–63). Consistent with our data, a preoperative study showed that mTORC1 inhibitor RAD001 decreased cell proliferation, particularly in aggressive and high-grade breast cancer patients (64).

CBX2 and CBX7 play oncogenic and tumor-suppressive roles, respectively, in breast cancer as reported earlier (34, 36). This study provides a mechanistic basis for such roles, particularly from a metabolic perspective, thus, imparting functional relevance to the biology of CBX2/7 in breast cancer. Glucose addiction represents a metabolic vulnerability and gaining insights into its regulation in cancer may provide important clues that could be exploited for therapeutic benefit. In the past decade, a wealth of knowledge understanding the aberrant glycolysis in cancer has been generated, yet, clinical targeting of glycolysis is far from giving desired results. This underlines the need for better understanding of glycolysis regulation in cancer. This work attempts to address some of these issues, at least in part, by identifying that CBX2/7-driven aerobic glycolysis is associated with breast tumor aggressiveness and poor prognosis. Based on the multi-omics data from breast tumors along with *in vitro* substantiation, we make a case for targeting of CBX2/7 and/or metabolic reprogramming in breast cancer for improved patient outcome.

Aggressive breast tumors are hard to treat and exhibit drug resistance, therefore, the patient outcome is poor despite substantial advancements in breast cancer therapeutics. There is a need to identify oncogenic processes driving aggressive breast cancer. The strong association of CBX2 and glycolysis deregulation with breast cancer aggressiveness suggests the therapeutic potential of targeting oncogenic CBX2 and/or glycolysis. As altered glycolysis has been suggested as a contributor to drug resistance in breast cancer (65–68), results of this work may have implications in strategies targeting glycolysis to overcome clinical drug resistance. Further, variation in patients response to chemotherapeutic drugs is a major obstacle faced in clinics. Drug sensitivity analysis demonstrating the utility of CBX2 expression in predicting sensitivity to methotrexate, rapamycin is a clinically relevant finding which may help in estimating the likelihood of patients benefitting from treatment of these drugs. Altogether, the results of this work postulate that CBX2-driven metabolic reprogramming may be a target-of-interest for aggressive breast cancer therapy.

The data presented signify the need of a better understanding of CBX2/7 biology and, invite further research into the how these readers of same histone code (trimethylation of lysine residue of H3) differentially alter metabolic reprogramming in breast cancer. A deeper mechanistic insight is needed to unravel whether CBX2/7 function independent of its epigenetic role to bring about metabolic changes in breast cancer. Additionally, results also shed light on the plausible role of CBX/PRC1 in metabolic reprogramming during embryonic development. Regardless, CBX2/7 expression status informs about the metabolic phenotype of breast tumors, patient outcome and sensitivity to anti-cancer drugs with potential implications in current or future anti-cancer strategies targeting clinical breast tumors.

## 1.4 Materials and methods

### 1.4.1 Transcriptomic, metabolomic and survival analysis of breast cancer patients

METABRIC breast tumor and normals data of 1992 and 144 samples, respectively, were accessed from European-Genome Archive (EGA) with accession numbers-EGAD00010000210, EGAD00010000211 and EGAD00010000212. TCGA breast tumor and normal data of 1104 and 102 samples respectively were obtained from UCSC Xena (https://xena.ucsc.edu). Metabolomics and transcriptomics data of 67 breast tumors and 65 normals was obtained from Terunuma et al. (41). DNA methylation and phosphoprotein data of TCGA tumor samples was obtained from cBioportal. Pathifier tool (https://www.bioconductor.org/packages/release/bioc/html/pathifier.html) was used in R script to calculate deregulation score of the studied pathway(s) in each tumor sample. Pathifier quantifies and assigns pathway deregulation scores (PDS) to each tumor sample based on gene-expression data and final PDS values are normalized between 0 and 1. A PDS value represents the extent of deviation of a pathway in a tumor sample from normal behavior. Normal tissue samples are required by Pathifier as a reference for normal gene-expression and to estimate deviation in terms of PDS. Signaling genesets used for Pathifier analysis (40) were taken from Molecular Signature Databse (MSigDB) (45) and glycolysis from Recon 1 (39). For patient survival analysis, Kaplan–Meier curves were prepared; *P* values, hazard ratios (HR) were calculated using the Mantel–Cox and log-rank test respectively, through GraphPad Prism software v7. For survival analysis, data was obtained from cBioportal.

### 1.4.2 Cell culture-based assays, siRNA transfections and Western blotting

MDA-MB-231 and MCF7 breast cancer cell lines were procured and maintained as described previously (21). Briefly, cells were grown in DMEM media (Gibco, ThermoFisher Scientific Inc., Waltham, MA, USA) supplemented with 10% FBS (Gibco) as described (21). Cell lines were authenticated using GenePrint system (Promega, WI, USA) and mycoplasma testing was performed using commercial detection kit (Thermo Fisher Scientific Inc.), to ensure authenticity and mycoplasma negativity. For CBX2 and CBX7 silencing experiments, MDA-MB-231 and MCF7 were seeded at a density of 1×10^4^ and 5×10^3^ per wells, respectively, followed by transfection with siRNA SMARTpool and non-targeting pool siRNA (Dharmacon, Lafayette, CO, USA). Pooled siRNA used combines four gene-specific siRNA into a single reagent pool. siRNA preparations and transfections were done according to the manufacturer’s protocol. Briefly, siRNAs were resuspended in 1 x siRNA buffer and cells were transfected using DharmaFECT transfection reagent (Dharmacon) according to manufacturer’s specifications. Cells were allowed to grow for 48 hours before processing and harvesting for experimental measurements. Glucose, lactate and ATP measurements were taken spectrophotometrically using commercial kits as described previously (21). Oligomycin and rapamycin (Sigma-Aldrich, St. Louis, MO, USA) were dissolved in cell-culture grade DMSO (Sigma-Aldrich) to prepare a 5 mM and 10 mM stock and solution and stored at −80°C until further use. All metabolic measurements were normalized to protein content. Cell viability and biomass experiments were performed using trypan blue exclusion and biomass measurements were performed using sulforhodamine-based (SRB) assays as described (21). For proliferation assay, cells were counted using a hemocytometer. For protein detection: cell lysates were prepared in RIPA lysis buffer containing protease and phosphates inhibitors (Sigma-Aldrich) and Western blotting was performed as described (69). Briefly, cells were incubated in lysis buffer for 30 min at 4°C with continuous shaking and then centrifuged to collect clear supernatant for protein quantification was done using Pierce™ BCA protein assay kit (Thermo Fisher Scientific Inc.). Primary antibodies used: anti-CBX2 (Abcam, Cambridge, UK), anti-CBX7 (Abcam), anti-phosphoS6-S235/236 (Cell Signaling Technologies, Danvers, MA, USA), anti-phosphoS6-S240/244 (Cell Signaling Technologies), anti-β-actin (Cell Signaling Technologies). The membrane was incubated with secondary antibody for 1 hour at room temperature and proteins were detected using chemiluminescent HRP substrate (Merck-Millipore, Merck KGaA, Darmstadt, Germany).

### 1.4.3 Drug sensitivity assay

Breast cancer cell lines sensitivity data was obtained from Heiser et al. (48). Expression data on breast cancer cell lines were obtained from (ArrayExpress E-MTAB-181) (48). Above mean expression cell lines were labelled as CBX2/CBX7High and below mean expression as CBX2/ CBX7Low. Z-scores of −log_10_(GI_50_) values were used to plot heatmap using Morpheus software from Broad Institute (https://software.broadinstitute.org/morpheus/).

### 1.4.4 Statistical analysis

Data are presented as either as median with minimum and maximum values or mean ± SD. Unpaired student’s t-test or Kruskal-Wallis or ANOVA with multiple comparison test was performed using GraphPad Prism software v7 to calculate significance. *P*<0.05 was considered to be statistically significant and represented in figures as \**P*<0.03, \*\**P*<0.0021, *\*\*\*P* <0.0002, \*\*\*\**P*<0.0001. All experiments were performed in replicates and repeated independently at least three times to calculate significance.

## 1.5 Acknowledgements

Department of Science and Technology (DST), Govt. of India, for DST-INSPIRE faculty award and research grant to MAI. AUR acknowledges Science and Engineering Research Board (SERB), Govt. of India, for National Postdoctoral Fellowship (NPDF).

## 1.8 Author contributions

Conception and design: MAI; Data acquisition: MAI, SS, AUR, PS; Data analysis and interpretation: MAI, SS, FAS, AUR, DS; Manuscript writing: MAI; Manuscript revision: all authors; Final approval for submission: all authors

## 1.9 Data accessibility

Breast tumor data used is publicly available from European Genome Archive (EGA), UCSC Xena and cBioportal. METABRIC data accession numbers-EGAD00010000210, EGAD00010000211 and EGAD00010000212. TCGA data obtained from UCSC Xena. Survival analysis data obtained from cBioportal. Drug sensitivity data accessed from Heiser et al (Ref. 48) and expression data from ArrayExpress E-MTAB-181.

## Supplementary figures

**Fig. S1:**
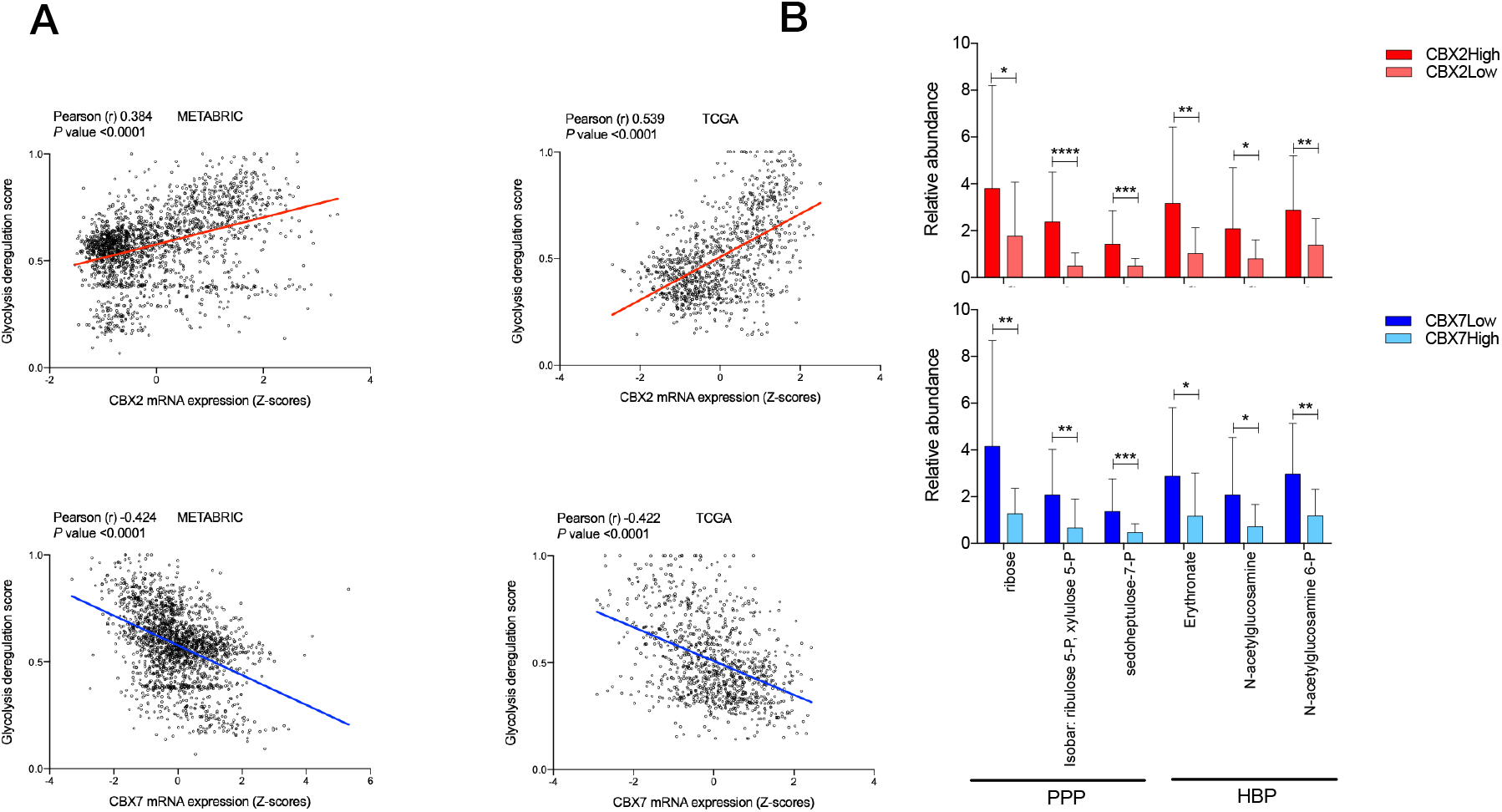
(A) Correlation plots showing positive and negative correlation of CBX2 and CBX7 with glycolysis PDS, respectively in METABRIC and TCGA. (B) Upregulated levels of key PPP and HBP metabolites in CBX2High and CBX7Low breast tumors, data from Terunuma et al (1). Data presented as mean ± SD. *P* values were calculated using *t*-test and represented as \**P*<0.03 and \*\**P<0.0021, ***P* <0.0002 and \*\*\*\**P*<0.0001.

**Fig. S2:**
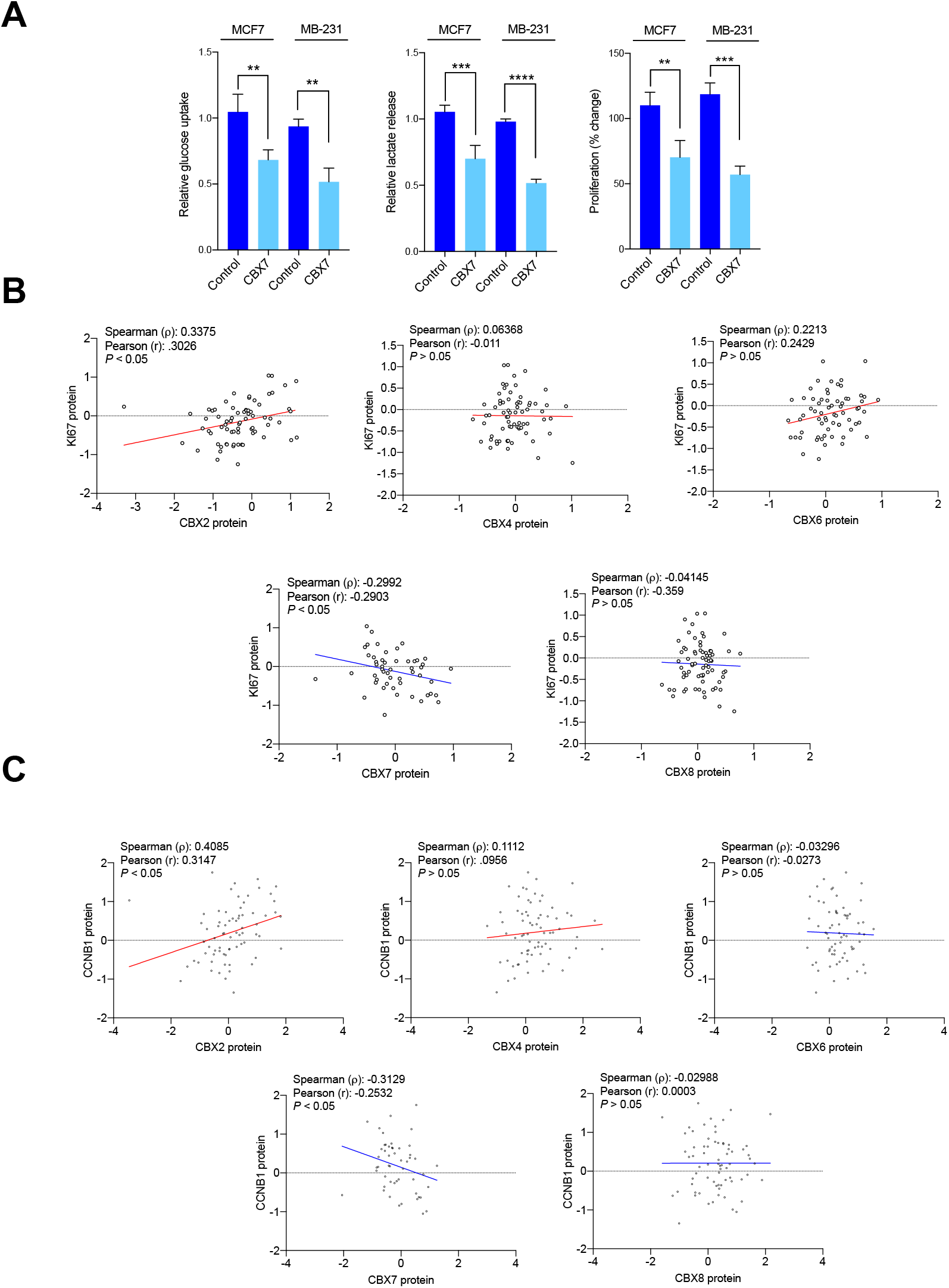
(A) Effect of CBX7 over-expression on glycolysis and proliferation. Correlation plots showing correlation of CBX2/4/6/7/8 with proliferation markers Ki67 and CCNB in TCGA breast tumors. Only CBX2/7 showed significant correlations in opposite directions. *P* values were calculated using Ptest and represented as \*\**P*<0.0021, *\*\*\*P* <0.0002, \*\*\*\**P*<0.0001.

**Fig. S3:**
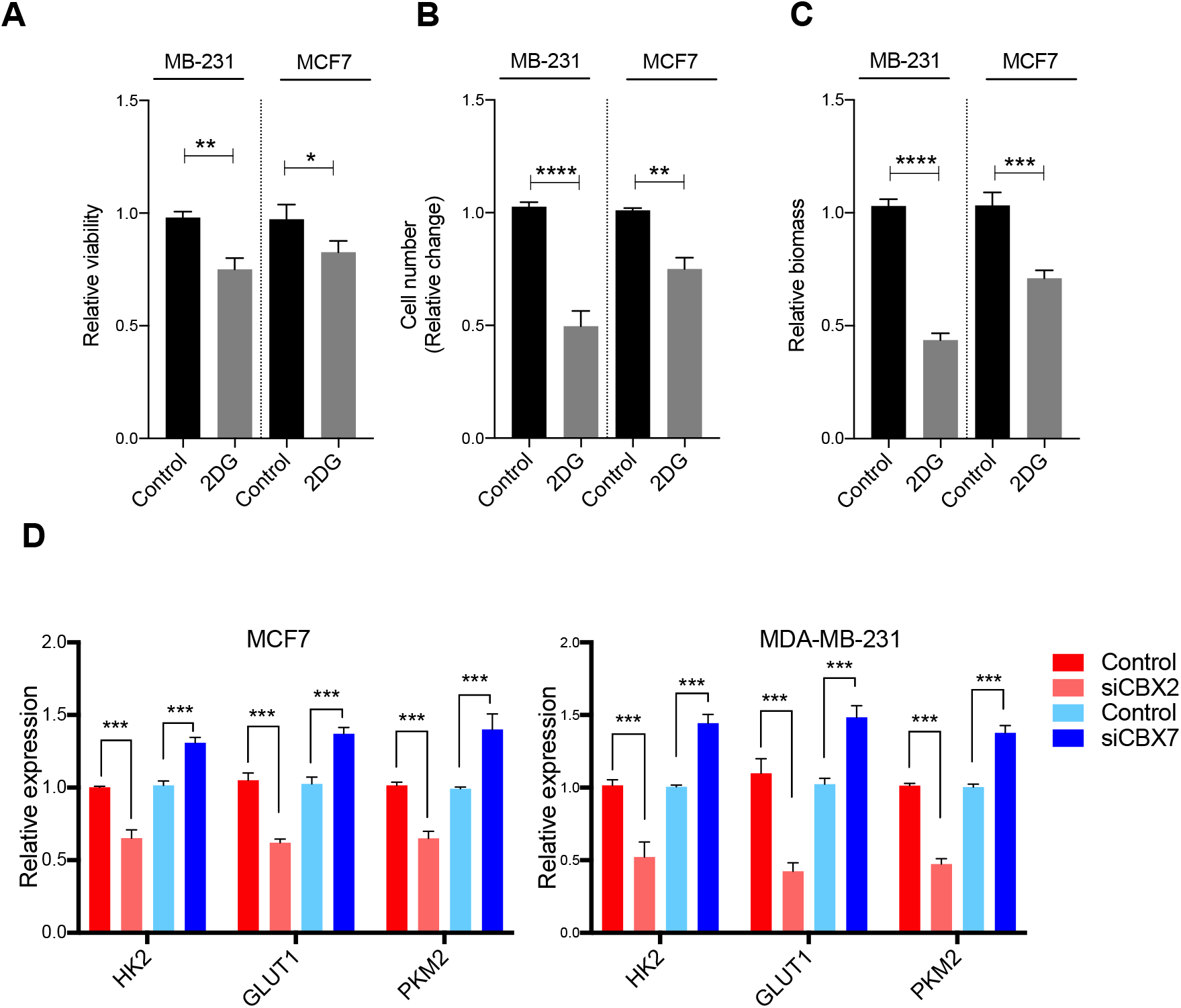
Effect of 10 mM 2-deoxyglucose (2DG) on (A) viability, (B) proliferation and (C) biomass of MDA-MB-231 and MCF7 cell lines after 48 hours of treatment. (D) Effect of CBX2/7 silencing on key glycolysis genes expression. Bars represent mean ± SD from independent experiments. *P* values were calculated using *t*-test and represented as \*\**P*<0.0021, \*\*\**P* <0.0002, \*\*\*\**P*<0.0001.

**Fig. S4:**
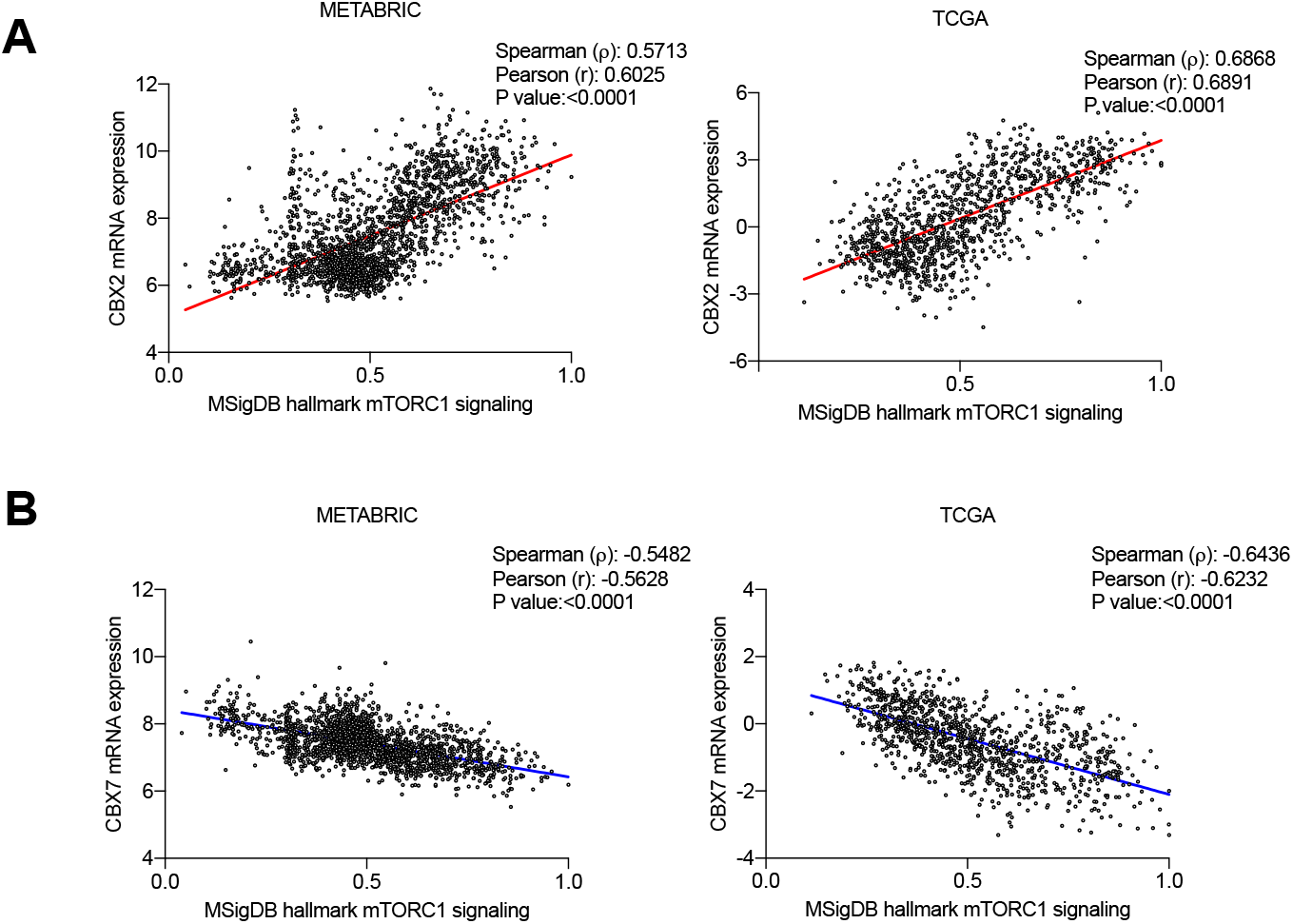
Correlation plots showing (A) positive and (B) negative correlations between CBX2 and CBX7, respectively, in METABRIC and TCGA datasets.

**Fig. S5:**
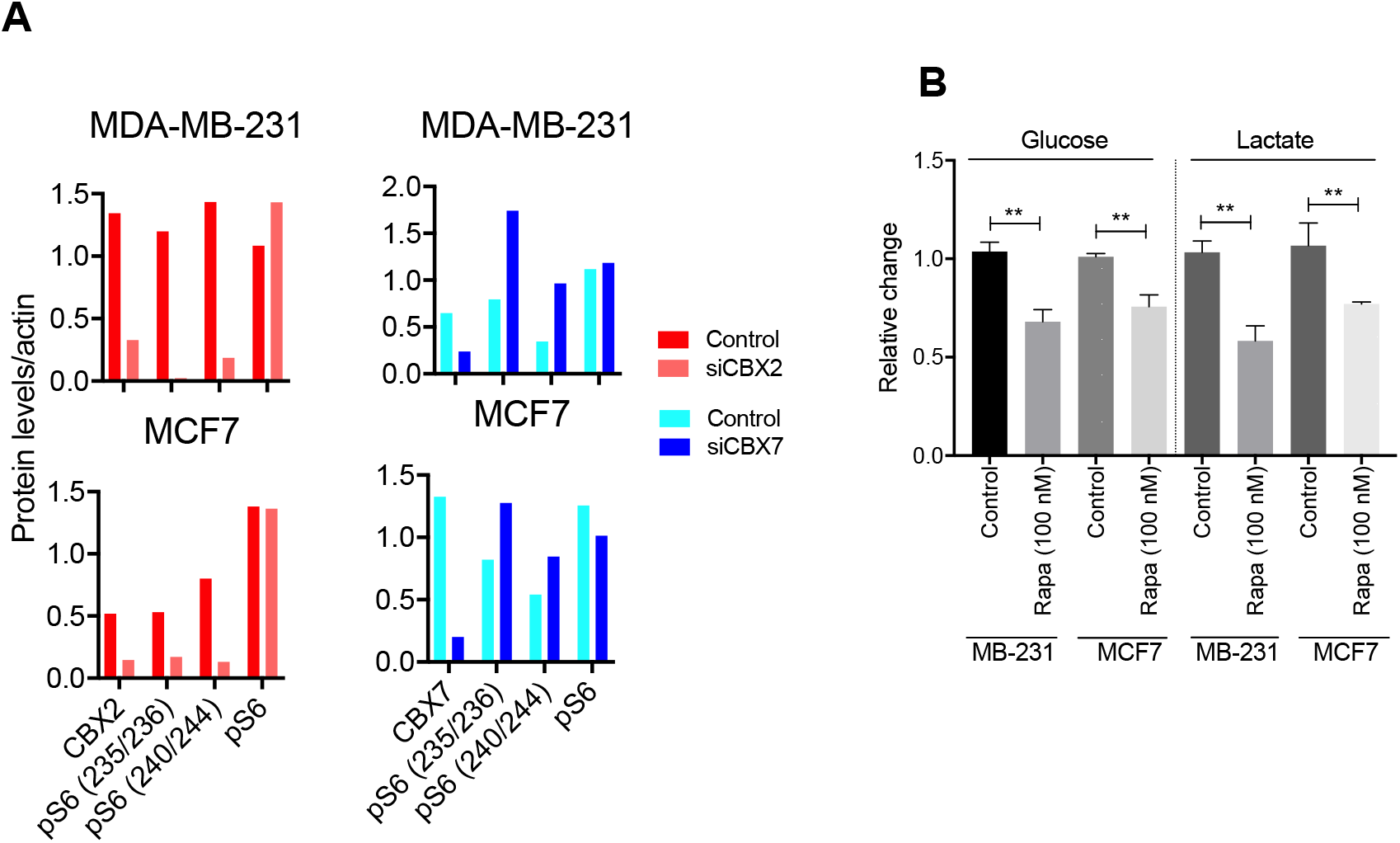
(A) Representative densitometric analysis related to Fig. 3. (B) Effect of rapamycin on glucose uptake and lactate in MDA-MB-231 and MCF7. Bars represent mean ± SD from 3 independent experiments. *P* value calculated using t-test and represented as \*\**P*<0.0021.

**Fig. S6:**
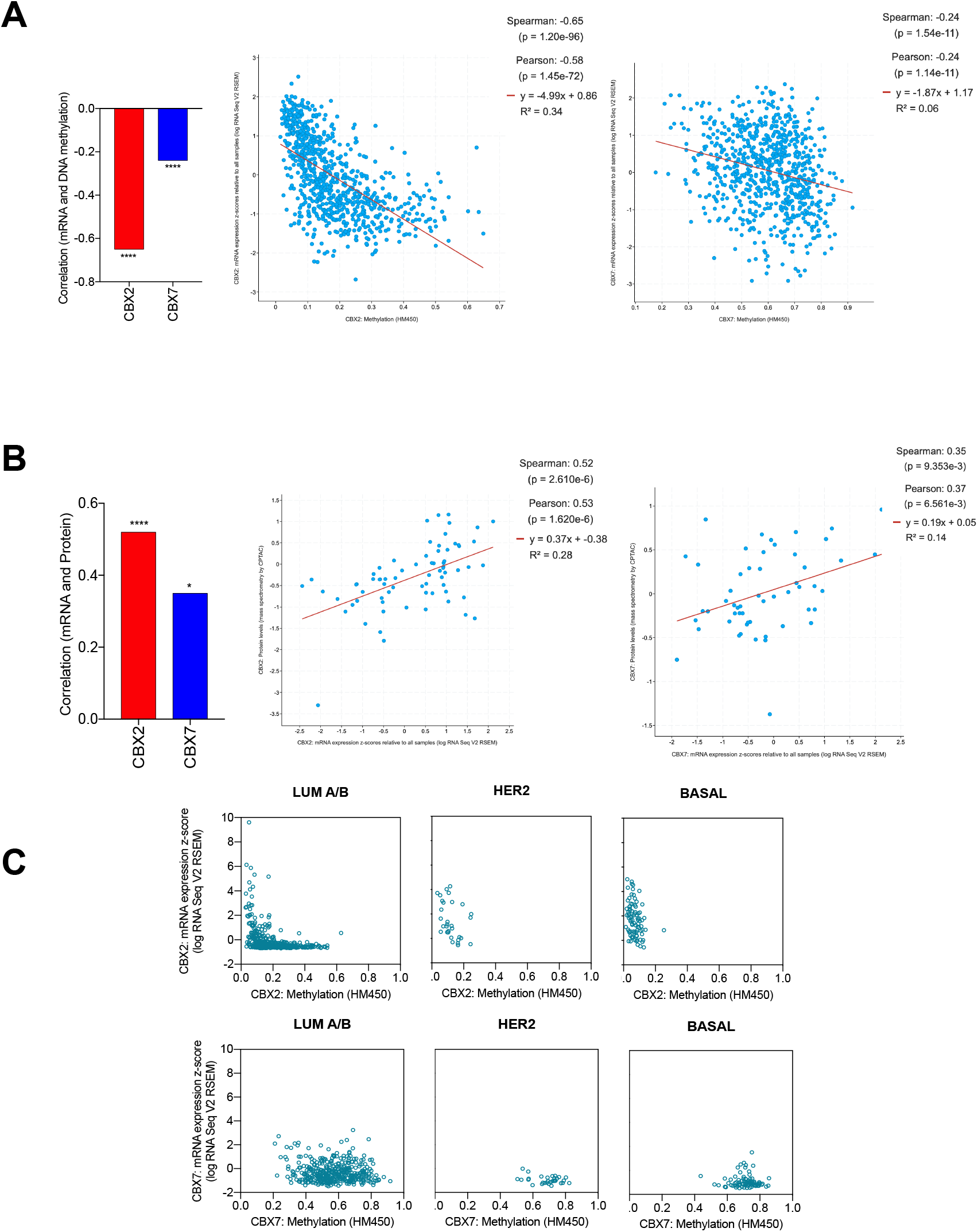
Negative correlation of CBX2 and CBX7 mRNA with (A) DNA methylation and positive correlation with (B) protein levels, in TCGA breast tumors. Subtype-specific correlation of CBX2/7 mRNA with DNA methylation. Data downloaded from cBioportal (2).

**Fig. S7:**
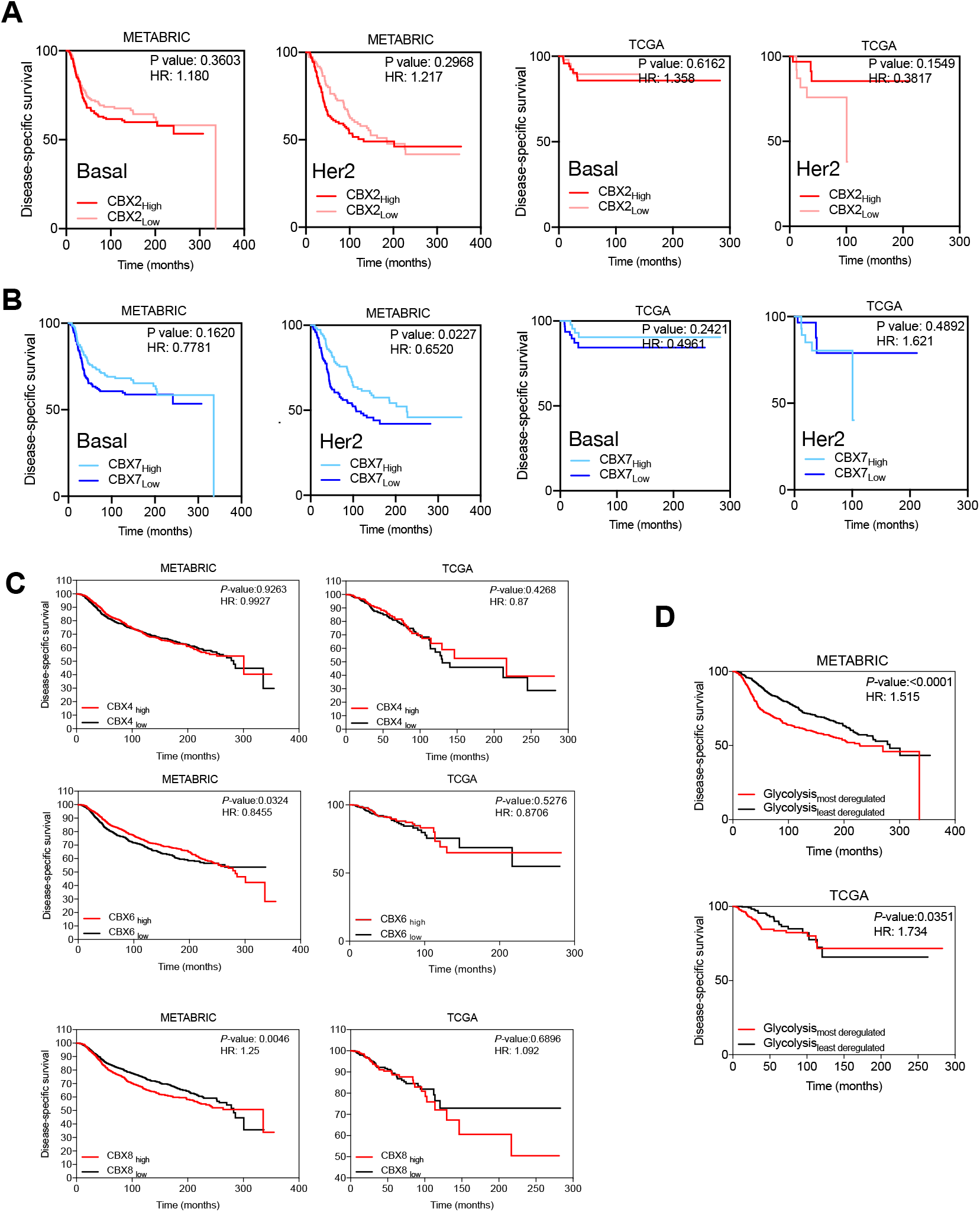
K-M curves showing prognostic relevance of (A) CBX2 and (B) CBX7 in basal and her2 samples. (C) CBX4/6/8 couldn’t predict prognosis reproducibly in both datasets. (D) Patients with higher deregulation of glycolysis showed poor outcome compared to patients with lower glycolysis deregulation scores.

**Fig S8:**
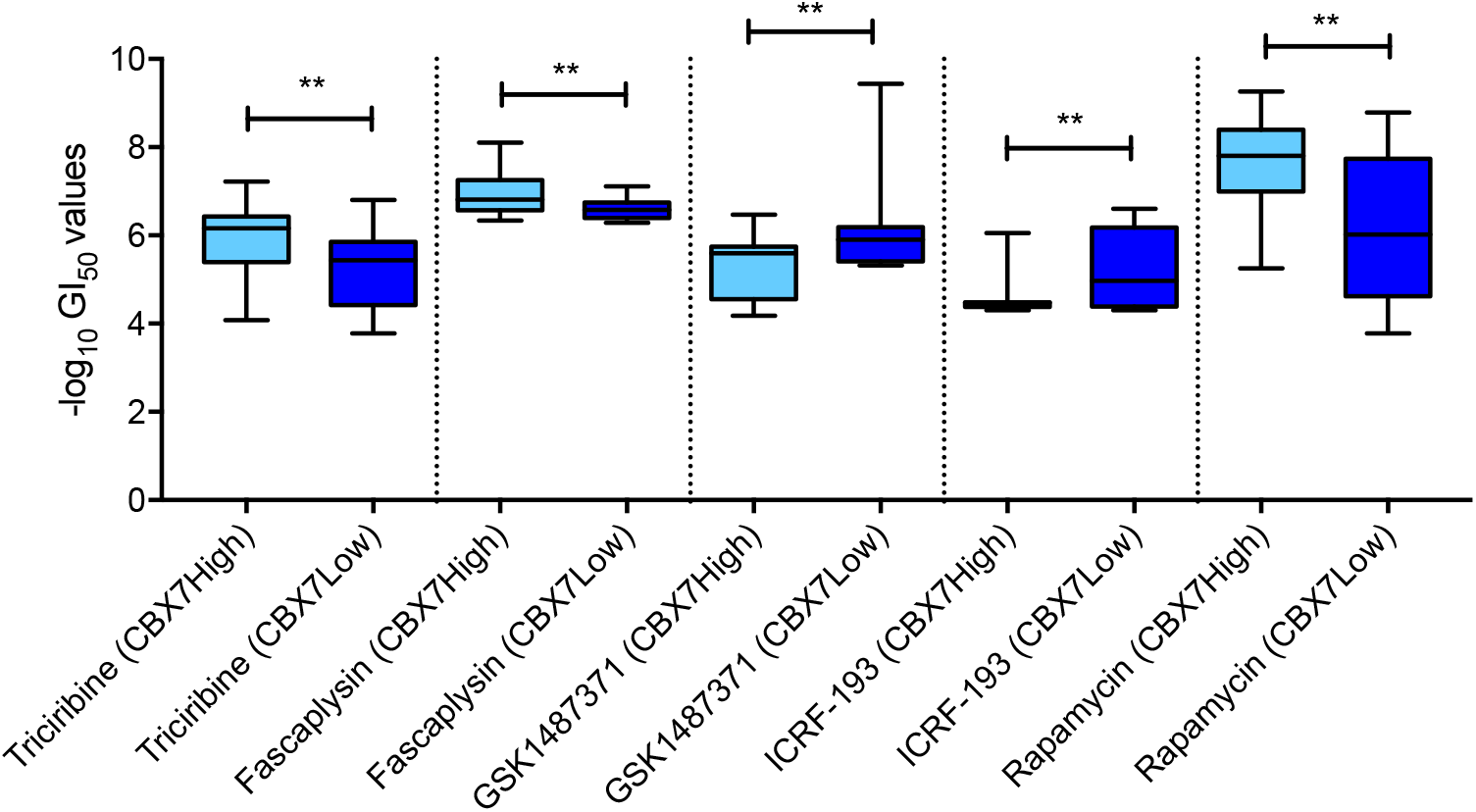
Difference is sensitivities to drugs in CBX7High/Low cell lines (see main text for details). Box and whiskers plot represent minimum, maximum and median. *P* values were calculated using *t*-test and represented as \**P*<0.03, \*\**P*<0.0021, *\*\*\*P* <0.0002, \*\*\*\**P*<0.0001.

