## Supplementary figures for "Multiomics integrative analysis reveals antagonistic roles of CBX2 and CBX7 in metabolic reprogramming of breast cancer"

### Supplementary figures 1-8

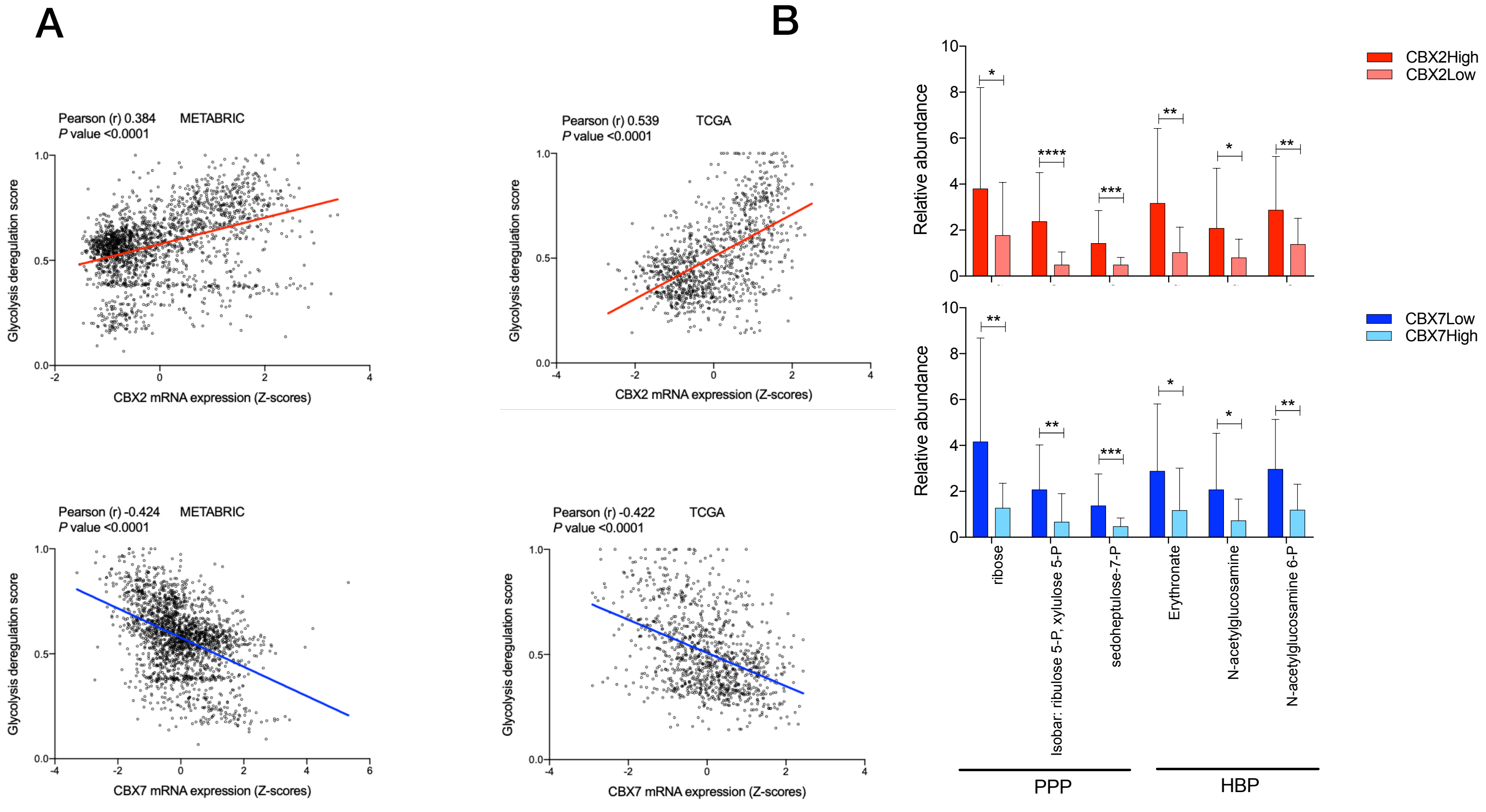


**Fig. S1:** (A) Correlation plots showing positive and negative correlation of CBX2 and CBX7 with glycolysis PDS, respectively in METABRIC and TCGA. (B) Upregulated levels of key PPP and HBP metabolites in CBX2High and CBX7Low breast tumors, data from Terunuma et al (1). Data presented as mean ± SD. *P* values were calculated using *t*-test and represented as **P*<0.03 and ***P*<0.0021, ****P* <0.0002 and *****P*<0.0001.

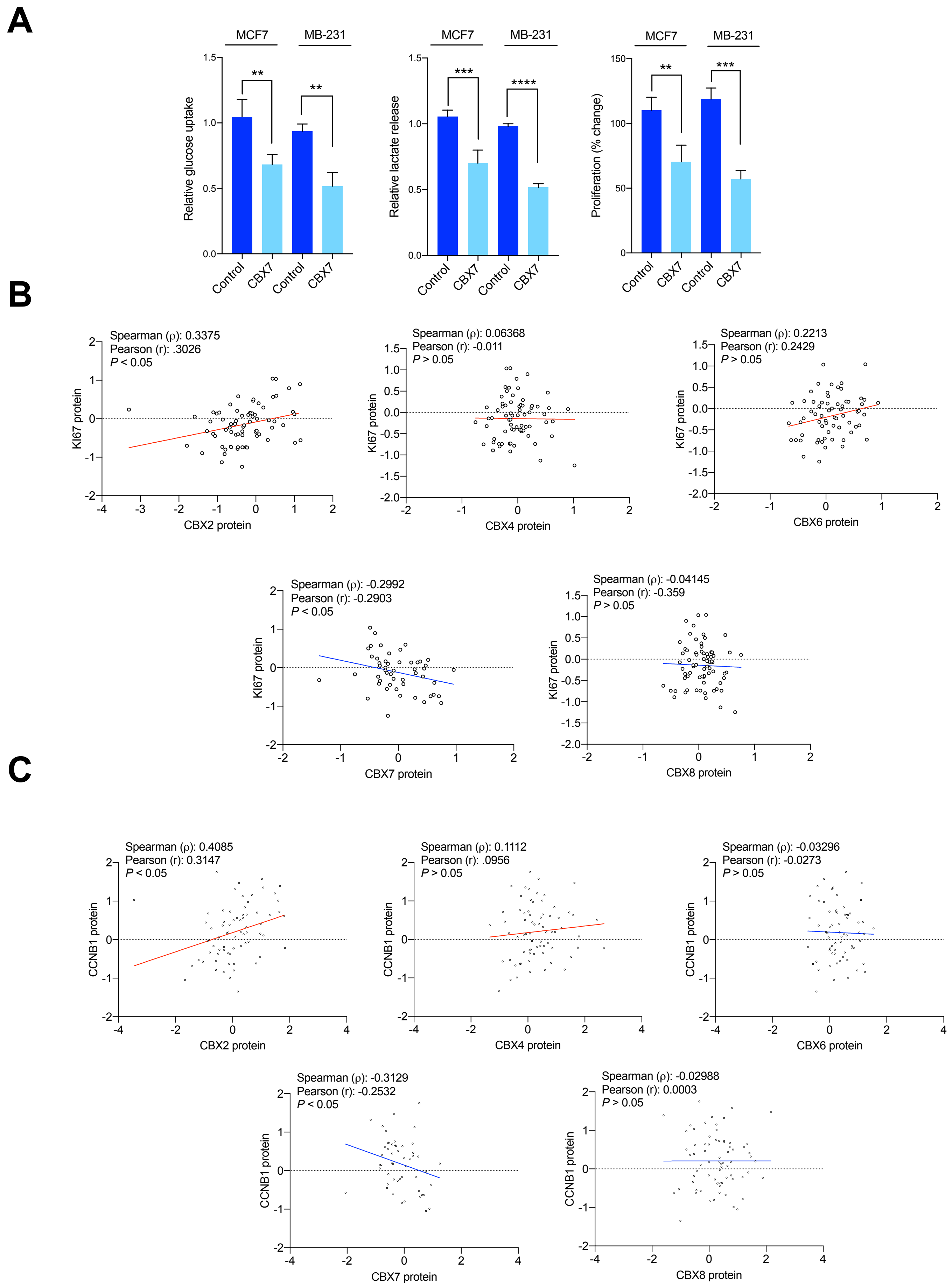


**Fig. S2:** (A) Effect of CBX7 over-expression on glycolysis and proliferation. Correlation plots showing correlation of CBX2/4/6/7/8 with proliferation markers Ki67 and CCNB in TCGA breast tumors. Only CBX2/7 showed significant correlations in opposite directions. *P* values were calculated using *t*-test and represented as $^{**}P<0.0021$, $^{***}P<0.0002$, $^{****}P<0.0001$.

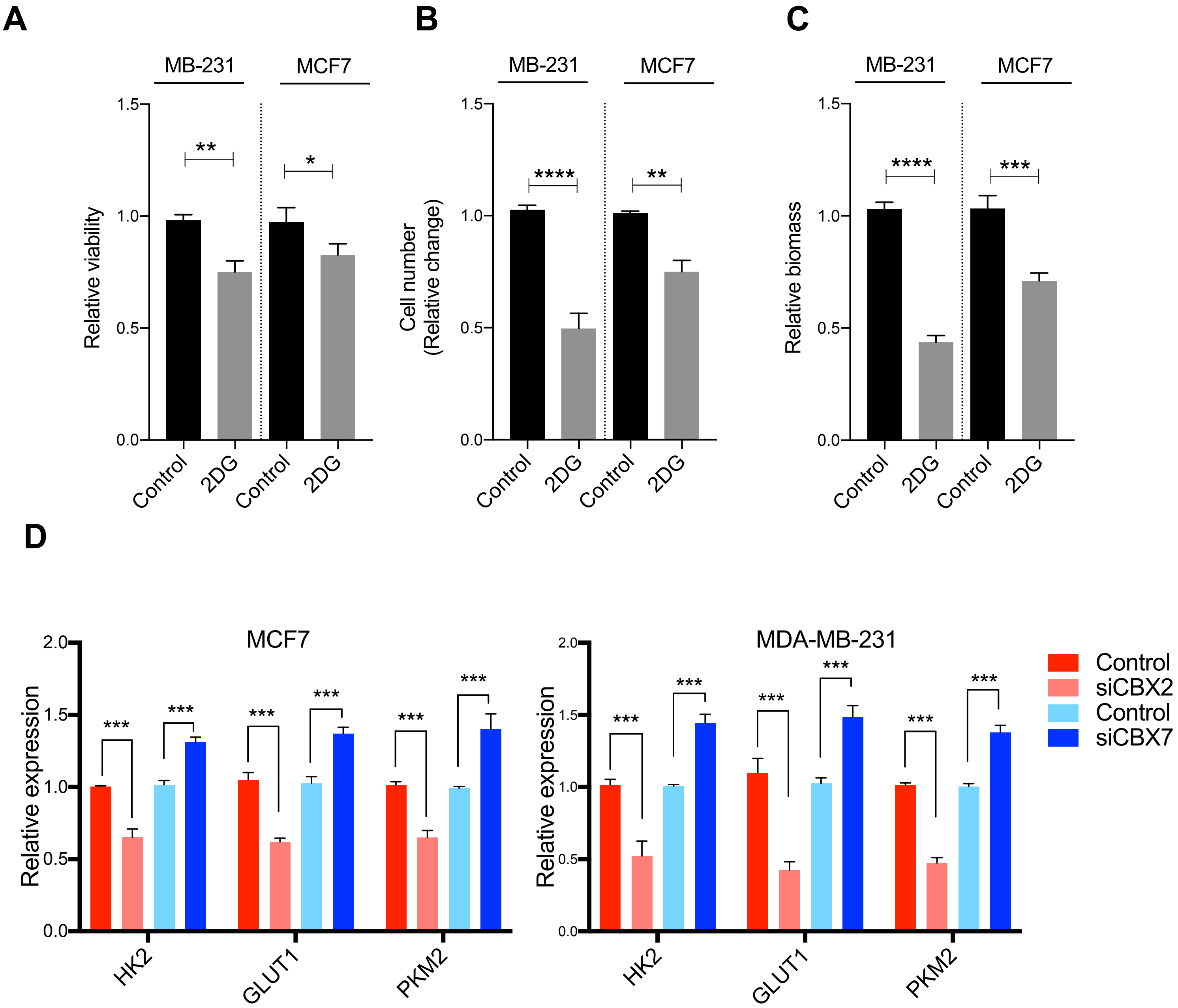


**Fig. S3:** Effect of 10 mM 2-deoxyglucose (2DG) on (A) viability, (B) proliferation and (C) biomass of MDA-MB-231 and MCF7 cell lines after 48 hours of treatment. (D) Effect of CBX2/7 silencing on key glycolysis genes expression. Bars represent mean ± SD from independent experiments. *P* values were calculated using *t*-test and represented as ***P*<0.0021, ****P* <0.0002, *****P*<0.0001.

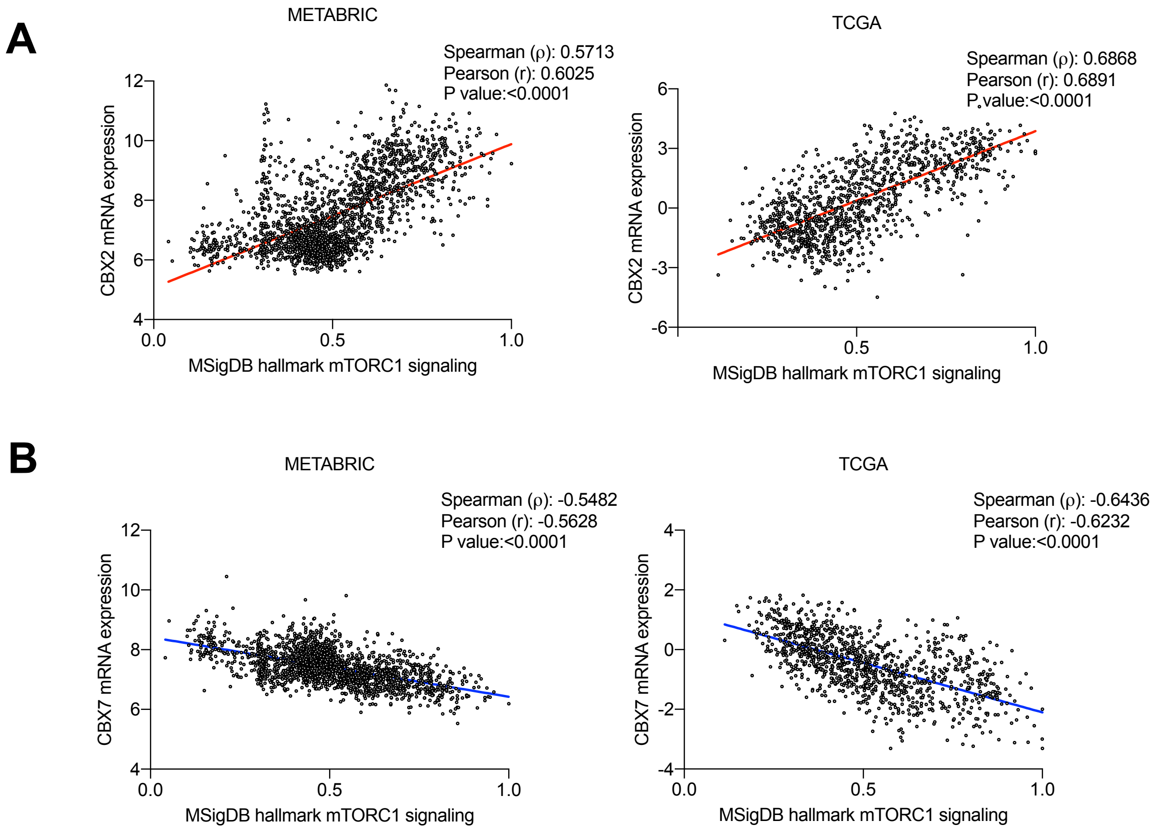


**Fig. S4:** Correlation plots showing (A) positive and (B) negative correlations between CBX2 and CBX7, respectively, in METABRIC and TCGA datasets.

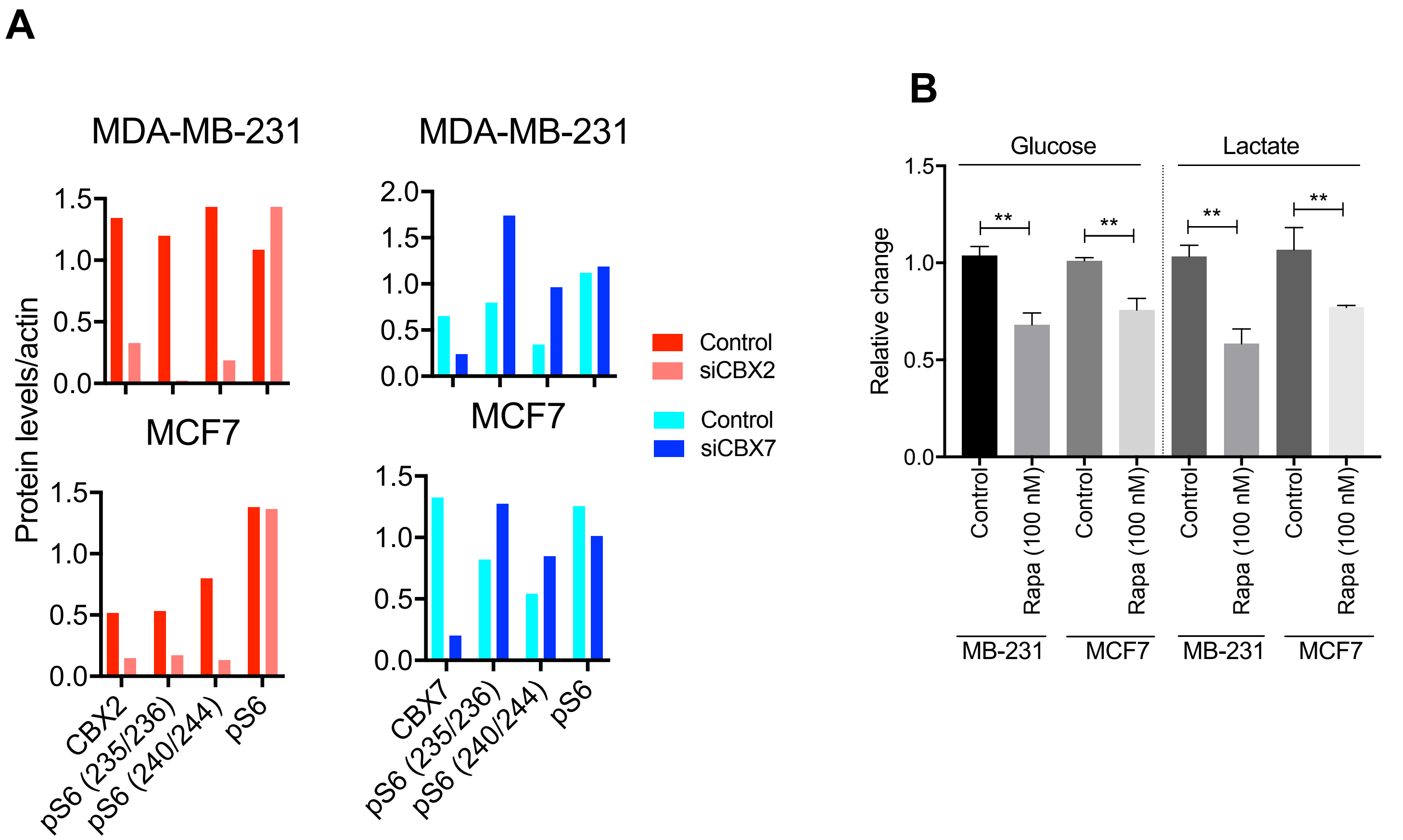


**Fig. S5:** (A) Representative densitometric analysis related to Fig. 3. (B) Effect of rapamycin on glucose uptake and lactate in MDA-MB-231 and MCF7. Bars represent mean ± SD from 3 independent experiments. *P* value calculated using t-test and represented as $^{**}P$<0.0021.

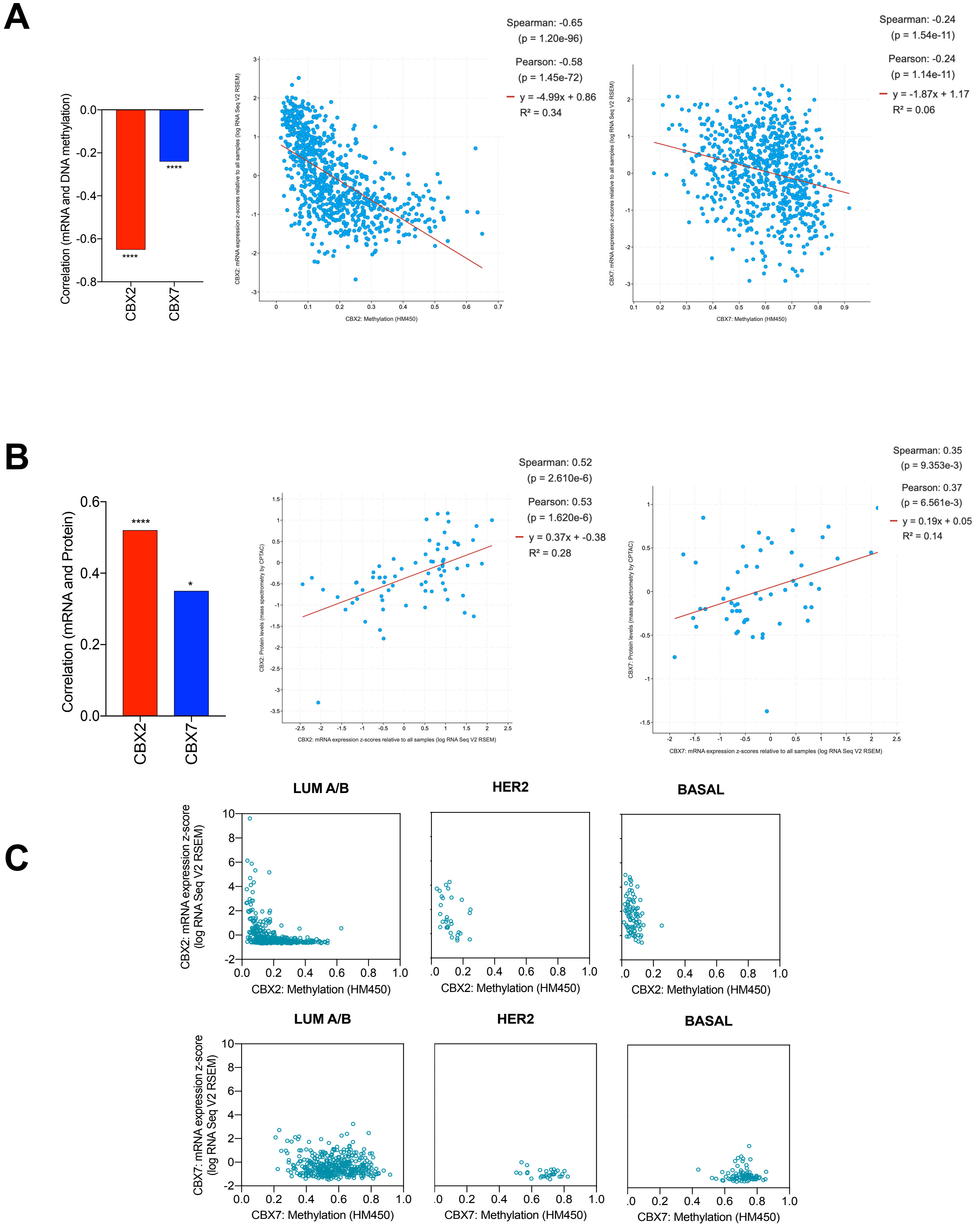


**Fig. S6:** Negative correlation of CBX2 and CBX7 mRNA with (A) DNA methylation and positive correlation with (B) protein levels, in TCGA breast tumors. Subtype-specific correlation of CBX2/7 mRNA with DNA methylation. Data downloaded from cBioportal (2).

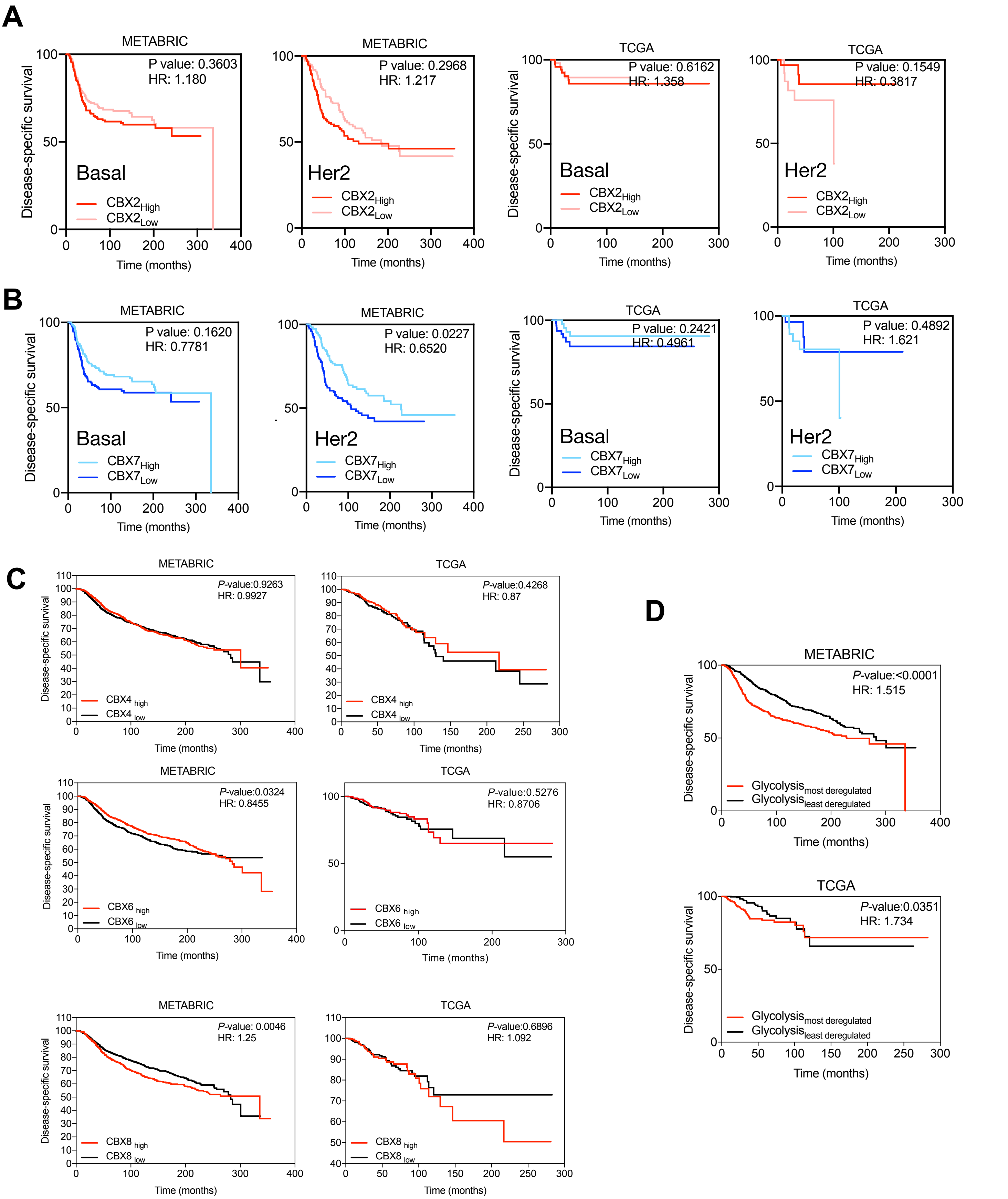


**Fig. S7:** K-M curves showing prognostic relevance of (A) CBX2 and (B) CBX7 in basal and her2 samples. (C) CBX4/6/8 couldn't predict prognosis reproducibly in both datasets. (D) Patients with higher deregulation of glycolysis showed poor outcome compared to patients with lower glycolysis deregulation scores.

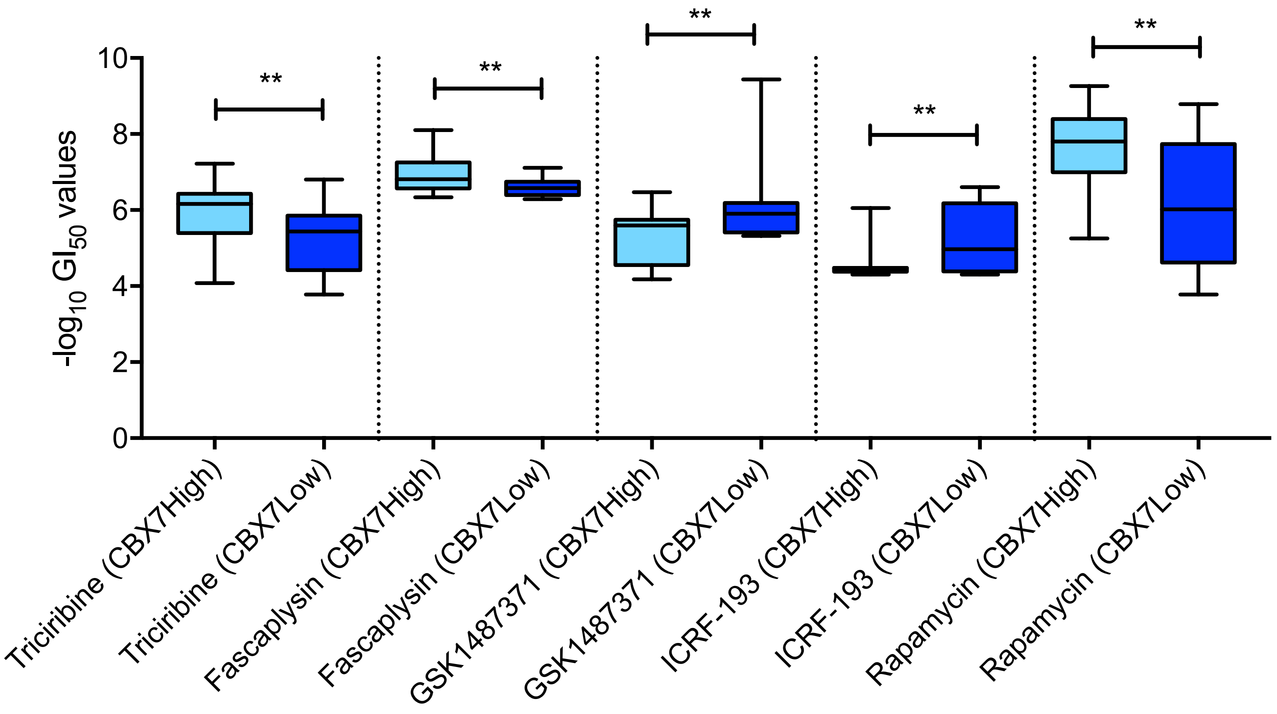


**Fig S8:** Difference is sensitivities to drugs in CBX7High/Low cell lines (see main text for details). Box and whiskers plot represent minimum, maximum and median. *P* values were calculated using *t*-test and represented as **P*<0.03, ***P*<0.0021, ****P* <0.0002, *****P*<0.0001.
